# Poorer Sleep Health is Associated With Altered Brain Activation During Cognitive Control Processing in Healthy Adults

**DOI:** 10.1101/2022.10.28.512671

**Authors:** Hanne Smevik, Sarah Habli, Simen Berg Saksvik, Elisabeth Kliem, Hallvard Røe Evensmoen, Virginia Conde, Agustin Petroni, Robert F. Asarnow, Emily L. Dennis, Live Eikenes, Håvard Kallestad, Trond Sand, Paul M. Thompson, Ingvild Saksvik-Lehouillier, Asta Kristine Håberg, Alexander Olsen

**Author notes:** Corresponding author: Alexander Olsen, PhD, Norwegian University of Science and Technology, Department of Psychology, 7491 Trondheim, Norway.

## Abstract

This study investigated how proactive and reactive cognitive control processing in the brain was associated with habitual sleep health. BOLD fMRI data was acquired from 81 healthy adults with normal sleep (41 females, age 20.96 - 39.58 years) during a test of cognitive control (Not-X CPT). Sleep health was assessed in the week before MRI scanning, using both objective (actigraphy) and self-report measures. Multiple measures indicating poorer sleep health - including later/more variable sleep timing, later chronotype preference, more insomnia symptoms and lower sleep efficiency - were associated with stronger and more widespread BOLD activations in fronto-parietal and subcortical brain regions during cognitive control processing (adjusted for age, sex, education, and fMRI task performance). Most associations were found for *reactive* cognitive control activation, indicating that poorer sleep health is linked to a ‘hyper-reactive’ brain state. Analysis of time-on-task effects showed that, with longer time on task, poorer sleep health was predominantly associated with increased *proactive* cognitive control activation, indicating recruitment of additional neural resources over time. Finally, shorter objective sleep duration was associated with lower BOLD activation with time on task and poorer task performance. In conclusion, even in ‘normal sleepers’, relatively poorer sleep health is associated with altered cognitive control processing, possibly reflecting compensatory mechanisms and / or inefficient neural processing.

---

Cognitive control underlies the regulation of thoughts, actions, and emotions, and relies on rapid, dynamic communication between widespread brain regions (Badre 2008; Braver 2012; Diamond 2013; Cole et al. 2014; Menon et al. 2020). Sleep is vital for brain health, everyday functioning, and quality of life (Walker and Stickgold 2006; Mignot 2008; Palmer and Alfano 2017; Tahmasian et al. 2020). Both cognitive control dysfunction and sleep-wake disturbances are transdiagnostic risk factors for developing mental health problems (McTeague et al. 2016; Freeman et al. 2020; Wainberg et al. 2021), and associated with negative outcomes across neurological and psychiatric disorders (Goschke 2013; Snyder et al. 2015). However, we still lack knowledge on potential links between cognitive control function and ‘normal’ habitual sleep in the general adult population, as most studies on this topic have focused on clinical, adolescent, or aging populations, and / or have included experimental manipulations of sleep or circadian rhythm (Scullin and Bliwise 2015; Krause et al. 2017; Lowe et al. 2017; Short et al. 2020; Tahmasian et al. 2021; Qin et al. 2023).

Even in ‘normal sleepers’ (persons without sleep or mental health complaints), there is considerable inter- and intraindividual variability in habitual sleep patterns and sleep need, i.e., sleep health (Buysse 2014; Beattie et al. 2015; Allen et al. 2018). Sleep health is a multidimensional construct - encompassing the duration, efficiency, timing, and subjective perception of sleep - and may be assessed via self-report (e.g., questionnaires) and objective (e.g., polysomnography, actigraphy) measures (Buysse 2014; van de Langenberg et al. 2022). Better sleep health is characterized by a regular sleep schedule with an appropriate sleep timing, adequate sleep duration (Hirshkowitz et al. 2015; Watson et al. 2015), good sleep efficiency (Ohayon et al. 2017), and experiencing little or no problems with sleep or daytime alertness (Buysse 2014; Allen et al. 2018). Meanwhile, indicators of poorer sleep health, including short sleep duration, poor sleep quality, and later chronotype (a preference for later sleep timing), have been linked with poorer health outcomes (Itani et al. 2017; Knutson and von Schantz 2018; Dong et al. 2019; Freeman et al. 2020).

To date, most studies on habitual sleep and cognitive function have relied on retrospective, self-report measures of singular aspects of sleep (typically sleep duration and/or quality) (Buysse 2014; Scullin and Bliwise 2015; van de Langenberg et al. 2022; Qin et al. 2023). This limits the interpretation of results, as self-report measures may be biased by a multitude of contextual and personal factors (Bliwise and Young 2007; Lauderdale et al. 2008; Lavie 2009; Matthews et al. 2018; Robbins et al. 2021). Furthermore, self-report and objective sleep measures - even within the same dimension - have low-to-modest correlations (Landry et al. 2015; Matthews et al. 2018; Thurman et al. 2018) and appear to be differently associated with cognitive function (Bernstein et al. 2019; McSorley et al. 2019; Hokett et al. 2021; Scarlett et al. 2021). While several large-scale studies have demonstrated an inverse U-shaped relationship between *self-reported* sleep duration and cognitive performance (poorer cognitive performance with both shorter and longer sleep durations) (Richards et al. 2017; Wild et al. 2018; Mantua and Simonelli 2019; Tai et al. 2022), studies using *objective* assessment of adult habitual sleep duration - which are far fewer in number - have provided mixed results (Scullin and Bliwise 2015; Fueggle et al. 2018; Kato et al. 2018; Scarlett et al. 2021; Suemoto et al. 2022; Qin et al. 2023). There is also growing evidence that other dimensions of habitual sleep health (such as the timing / variability and efficiency of sleep) are important for cognitive function (Owens et al. 2016; Facer-Childs et al. 2019; Hershner 2020; Zhang et al. 2020; Stefansdottir et al. 2022; Qin et al. 2023). However, these dimensions are more difficult to quantify (Buysse 2014), and the data on sleep variability are not presently viewed as robust (Bei et al. 2016; Chaput et al. 2020). To further elucidate the links between sleep and cognitive function, there is a need to include multidimensional assessment of ‘normal’, habitual sleep health, using both subjective and objective measures (Chaput et al. 2020; van de Langenberg et al. 2022).

The interplay between sleep and cognitive function is likely mediated by brain structure and functioning (Avinun et al. 2017; Alfini et al. 2020; Grumbach et al. 2020; Tahmasian et al. 2020; Schiel et al. 2022). Measures of poorer sleep health have been associated with lower white matter integrity (Yaffe et al. 2016; Khalsa et al. 2017; Grumbach et al. 2020), lower gray matter volume and thickness (Sexton et al. 2014; Cheng et al. 2020; Kim et al. 2021; Wang et al. 2021), as well as altered functional connectivity within and between widespread brain regions (Curtis et al. 2016; Cheng et al. 2018; Tashjian et al. 2018; Lunsford-Avery et al. 2020). In task-based functional magnetic resonance imaging (fMRI) studies, shorter habitual sleep has been associated with reduced frontal, occipital, and insular blood oxygen level-dependent (BOLD) activations during negative distractor processing (Dimitrov et al. 2021), threat perception (Tashjian and Galván 2020), and risky decision making under stress (Uy and Galván 2017). Poorer self-reported sleep quality has been linked to lower activation within the insular and anterior cingulate cortices during emotion processing (Klumpp et al. 2017; Guadagni et al. 2018), and inconsistent sleep timing has been linked to less occipital activation and worse task performance during high working-memory loads (Zhang et al. 2020). In sum, the emerging pattern is that different indicators of poorer sleep health are associated with altered functional and structural characteristics of specific brain regions important for cognitive control function (Dosenbach et al. 2008; Olsen et al. 2013; Cole et al. 2014). However, there is a lack of theoretical frameworks to aid interpretation of extant literature, which is based on heterogeneous study samples, and mainly comprises evidence from structural and resting-state neuroimaging methods, as well as task-based activity related to affective processing (Salehinejad et al. 2021).

The *dual mechanisms framework of cognitive control* focuses on the temporal aspects of cognitive control processing (Braver 2012). In this framework, *proactive* cognitive control supports processing occurring over relatively longer time periods, such as the maintenance of goal-relevant information and continuous monitoring of incoming stimuli. *Reactive* cognitive control underlies more rapid processing occurring on a trial-to-trial basis, and typically engages when a conflicting or interfering stimulus is detected. Anatomically, these temporal modes of cognitive control rely on distinct, yet closely interacting brain networks (Seeley et al. 2007; Dosenbach et al. 2008; Braver 2012; Olsen et al. 2013; Cole et al. 2014), and converge on “core” control regions including fronto-parietal, insular, and subcortical areas (Dosenbach et al. 2006; Niendam et al. 2012; Olsen et al. 2013; Cole et al. 2014; Cai et al. 2016). Optimal cognitive functioning relies on a dynamic interplay between proactive and reactive cognitive control processes (Braver 2012; Olsen et al. 2013), where either type of processing may be more or less beneficial depending on context (Lesh et al. 2013; Vanderhasselt et al. 2014; Olsen et al. 2015, 2018; Kleerekooper et al. 2016; Huang et al. 2017). A relative increase in proactive cognitive control processing is associated with healthy brain development (Staub et al. 2014; Chevalier et al. 2015; Kubota et al. 2020; Niebaum et al. 2020), and has been linked to compensatory mechanisms associated with better every day cognitive control function in people with moderate/severe traumatic brain injury (Olsen et al. 2015). On the other hand, a relative increase in reactive cognitive control processing has been linked to poorer white matter organization, lower fluid intelligence, as well as higher levels of anxiety and stress (Fales et al. 2008; Burgess and Braver 2010; Schmid et al. 2015; Olsen et al. 2018; Husa et al. 2022).

Another temporal aspect of cognitive control function is the effect of time on task, typically observed as decline in performance and vigilance with sustained task performance (Langner and Eickhoff 2013). Such effects have been linked to mental fatigue and depletion of cognitive resources over time (Lim et al. 2010). Proactive cognitive control processing seems to be more sensitive to time-on-task effects (Olsen et al. 2013, 2015), possibly due to having a higher metabolic cost (engaging more wide-spread brain areas over longer time periods), and consequently being more taxing to uphold over time (Burgess and Braver 2010; Braver 2012). Importantly, time-on-task effects are exacerbated by sleep deprivation (Hudson et al. 2020), and the two have been linked to brain activity changes in overlapping areas, suggesting possibly shared underlying mechanisms (Asplund and Chee 2013; Satterfield et al. 2017). Investigating associations between time-on-task effects and habitual sleep health may therefore shed light on more subtle neural processes underlying the association between sleep and cognitive function in the healthy brain.

The main aim of this study was to investigate whether and how proactive and reactive cognitive control processing in the brain are associated with sleep health in adult, ‘normal sleepers’. To this end, we used a well-validated, fMRI-adapted continuous performance test which was specifically developed to assess the temporal dynamics of cognitive control, including time-on-task effects (Olsen et al. 2013, 2015). Our primary analysis (1) investigated associations between sleep health and BOLD activation during proactive and reactive cognitive control processing. As a secondary analysis (2), we examined the associations between sleep health and changes in BOLD activations (proactive and reactive cognitive control) with time on task. Age, sex, education, and fMRI task performance were selected as covariates of no interest a-priori, to control for possible confounding factors related to sleep health, cognitive control function, and BOLD activation (Price et al. 2006; Yarkoni et al. 2009; Olsen et al. 2015; Shanmugan and Satterthwaite 2016; Gurvich et al. 2018; Evans et al. 2021; van de Langenberg et al. 2022). This is the first study to investigate how sleep health may be associated with proactive and reactive cognitive control processing in the brain. We therefore took an *exploratory* approach aimed at capturing the multidimensionality of sleep health, and included a comprehensive selection of objective (actigraphy-based) and self-reported (questionnaire-based) measures, encompassing those measures which have been most often used in previous literature (e.g., Beattie et al. 2015; Klumpp et al. 2017; Grumbach et al. 2020; Zhang et al. 2020).

## Materials and Methods

### Study Design and Procedure

Data were collected as part of a randomized controlled trial spanning three weeks. The current study uses data from the first week of this trial (baseline - before randomization or any intervention). The study had a prospective design, which included two on-site visits separated by seven days (questionnaires and cognitive testing at visit 1, fMRI at visit 2; see Figure 1a). Naturalistic sleep-wake data were recorded between visits using actigraphy and sleep diaries. Participants were asked to ‘sleep as usual’ throughout the study period. At visit 1, participants performed a standardized test of cognitive control function and completed questionnaires assessing demographic information and sleep health. They received their actigraph and sleep diary and were instructed to keep the actigraphs on 24/7 throughout the study period.

**Figure 1.**
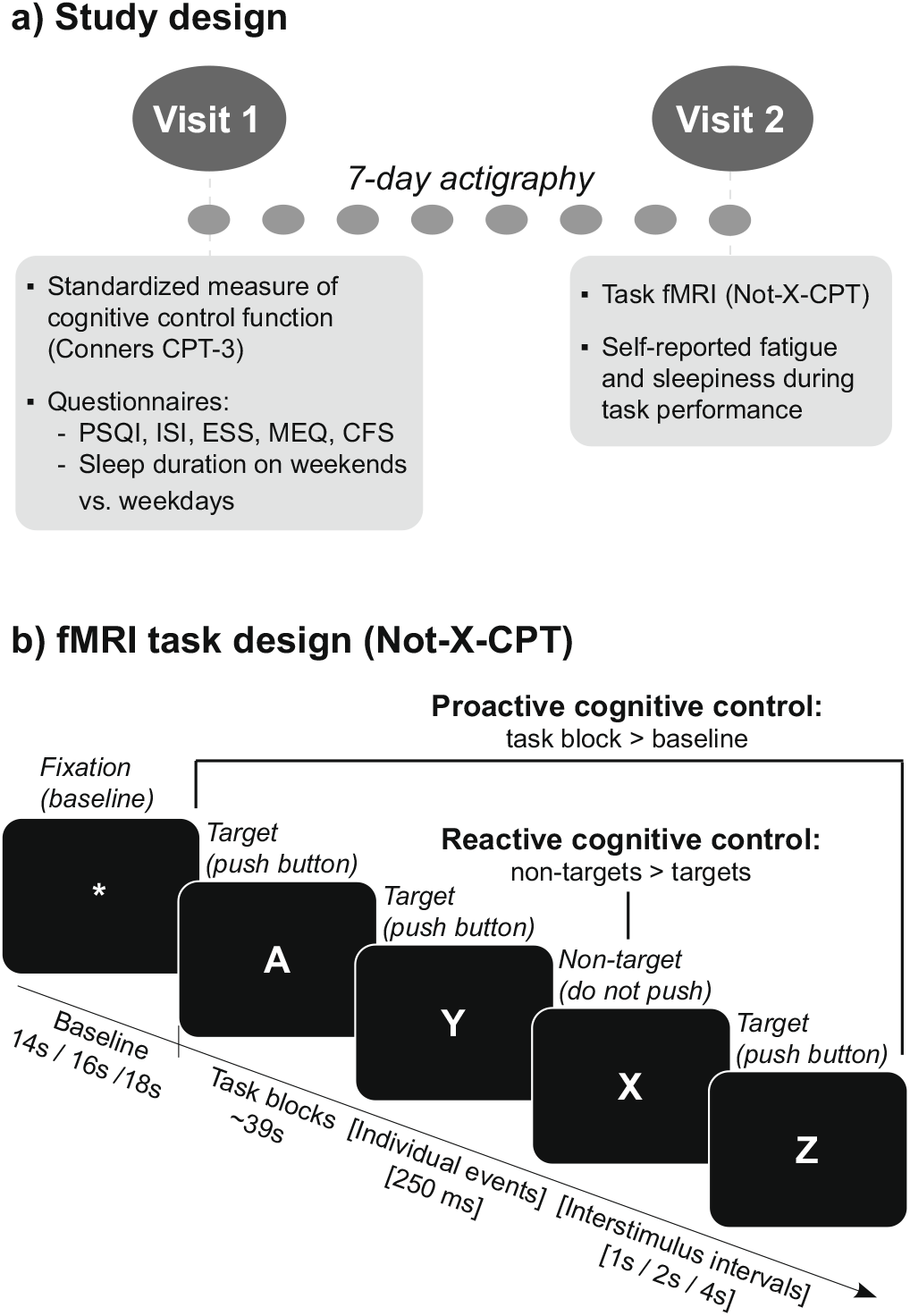
Overview of Study Design and Sleep Health Measures. a) Participants completed two study visits, and naturalistic, habitual sleep was measured during the 7 nights between visits using actigraphy. Visits were at the same time of day (between 8 AM and 3 PM) for each participant. At visit 1, participants completed a computerized, standardized test of cognitive control (Conners CPT-3) as well as a series of validated questionnaires on sleep health and fatigue. They also received actigraphs and sleep diaries (used for quality control of actigraphy data). At visit 2, participants completed a ~30 minute task fMRI session, and were asked to report on their current level of mental fatigue and sleepiness halfway through the task. b) To test cognitive control function, we used a ‘Not-X-CPT’ task adapted to a mixed block/event-related fMRI design (Olsen et al. 2013, 2018). Letters were consecutively presented on the screen and participants were asked to respond to press a response button as quickly and accurately as possible whenever a target (letters A-Z) was presented, and not respond when a non-target (letter X) was presented. The task consisted of a total of 480 stimuli (10% non-targets), with a stimulus duration of 250 milliseconds, and varying interstimulus intervals of 1, 2, or 4 seconds (jittered), to allow for event-related fMRI analysis (Petersen and Dubis 2012). The task was presented in two separate runs, each lasting ~15 minutes, containing 16 task blocks (duration ~39 seconds) and 16 baseline blocks (varying inter-block intervals of 14, 16 or 18 seconds). To eliminate systematic order effects, the different task parameters (inter-block intervals, stimulus type, block type and inter-stimulus intervals) were counterbalanced within and between the two task runs. See Olsen et al. (2013, 2018) for more details on the task design. For use in our primary/secondary analyses, the following contrasts were computed: (1) Proactive Cognitive Control (task blocks > fixations), Reactive Cognitive Control (non-targets > targets), as well as (2) time-on-task change for each contrast (Δ Proactive Cognitive Control and Δ Reactive Cognitive Control). CPT = Continuous Performance Test; MEQ = Morningness-Eveningness Questionnaire; PSQI = Pittsburgh Sleep quality Index; ISI = Insomnia Severity Index; ESS = Epworth Sleepiness Scale; CFS = Chalder Fatigue Scale; fMRI: functional magnetic resonance imaging.

At visit 2, participants completed a continuous performance test during fMRI (Not-X-CPT, Figure 1b) and were asked to report their levels of mental fatigue and sleepiness halfway through the task. All on-site testing was conducted between 8 AM and 3 PM. Participants were scheduled for testing at the same time of day on both visits to control for circadian effects in each individual. They were also asked to avoid caffeine and nicotine in the last two hours before testing.

### Participants

Healthy volunteers without sleep complaints (assessed using criteria described below) were recruited via online advertisements, public posters, and word-of-mouth in and around the city of Trondheim, Norway. Inclusion and exclusion criteria were assessed in a structured phone interview. Inclusion criteria were: 1) being between 20-40 years of age, 2) having normal or corrected-to-normal vision, and 3) fluency in the Norwegian language. Exclusion criteria were: 1) contraindications for MRI, 2) any chronic or ongoing medical, neurological or mental illness (including sleep disorders), and 3) obvious factor(s) that would likely influence results or adherence to the study protocol, such as night work or shift work, irregular sleep or perceived sleep problems, excessive alcohol use, previous habitual use of psychotropic drugs (e.g., marijuana, stimulants), or any use of such drugs in the last three months. Assessment of ‘normal sleep’ was based on the Research Diagnostic Criteria for Normal Sleepers (Edinger et al. 2004). According to these criteria the individuals must have (a) no complaints of sleep disturbance or daytime symptoms attributable to unsatisfactory sleep, (b) have a routine standard sleep/wake schedule characterized by regular bedtimes and rise times, (c) no evidence of a sleep-disruptive medical or mental disorder, (d) no evidence of sleep disruption due to a substance exposure, use, abuse or withdrawal, and (e) no evidence of a primary sleep disorder (Edinger et al. 2004). The criteria were operationalized and assessed in the structured interview at what is referred to as levels 1 & 2 (simple self-report and personal history) in recommendations made by Beattie et al. (2015). Importantly, ‘normal sleep’ is not necessarily equal to ‘good sleep’ (Buysse 2014; Beattie et al. 2015), and even in normal sleepers, there is an expected inter- and intraindividual variability in sleep health (sleep timing, duration and quality). Normal sleepers may therefore still report issues with sleep health as measured through a questionnaire (e.g., symptoms of insomnia) without meeting the criteria for a significant sleep complaint or clinical diagnosis (e.g., insomnia disorder).

A flowchart of the inclusion process is presented in Figure 2. Of the 92 participants enrolled in the study, four were excluded due to only having completed visit 1, four were excluded due to technical problems with the MRI scanner, one was excluded due to technical problems with logging task fMRI behavioral data, and two were excluded due to missing (n = 1) or poor (n = 1) actigraphy data. This left a final sample of 81 participants who completed both study visits and had usable fMRI, actigraphy and questionnaire data (41 women; mean age = 27.82 years, SD = 5.42; mean education = 16.28 years, SD = 2.34).

**Figure 2.**
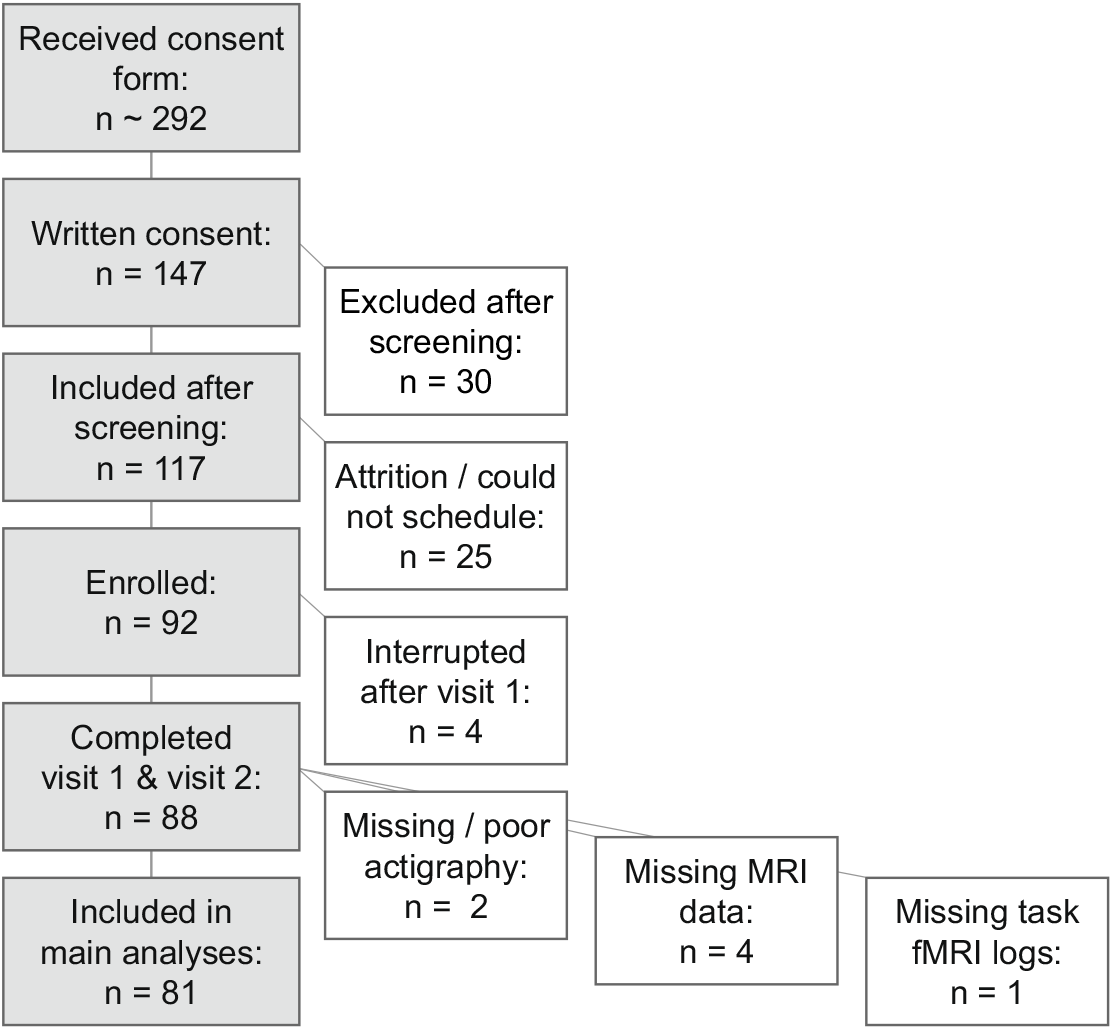
Overview of Inclusion Process. fMRI = functional magnetic resonance imaging

The study was approved by the Regional Committee for Medical and Health Research Ethics in Central Norway (REK number 2018/2413) and conducted in accordance with the 1964 Helsinki Declaration and its later amendments or comparable ethical standards.

### Measures of Sleep Health, Fatigue, and Standardized Measure of Cognitive Control Function

#### Objective Measures of Sleep Health

Objective, prospective assessment of participants’ habitual sleep was obtained using actigraphy (Actiwatch Spectrum Pro, Philips Respironics Inc., Murrysville, USA). The actigraphs recorded the participants’ light exposure and daily activity via a light sensor and a piezoelectric accelerometer, respectively. Data were recorded using 15-second epoch lengths. Participants were asked to use an event marker on the actigraph to indicate when they lay down to sleep each night. Sleep-wake periods were automatically classified using the Actiware software (Philips Actiware 6.0.9) using a medium sensitivity setting, and thereafter systematically inspected by trained study staff (Boyne et al. 2013). For qualitative assessment of the actigraphy data, participants filled out sleep diaries based on the Consensus sleep diary each morning (Carney et al. 2012). In cases of obvious misclassifications of rest periods, or large discrepancies between actigraphy and sleep diary data, rest interval onsets and offsets were manually adjusted according to event markers, activity, light levels, and / or sleep diaries using a standardized procedure (Follesø et al. 2021).

For each night, the following variables were extracted from the raw actigraphy data: sleep onset, sleep offset, sleep duration (total sleep time during the night, excluding brief wakes after sleep onset (WASO)), sleep efficiency (sleep duration divided by time in bed) and sleep onset latency (interval between going to bed and sleep onset (SOL)). Sleep midpoints were calculated by dividing the sleep interval (sleep offset - sleep onset) by 2 and subtracting the resulting duration from the sleep offset time point. Individual, 7-day averages of sleep duration, midpoint, efficiency, and SOL were calculated for use in statistical analyses, and the standard deviation of sleep duration and midpoint were used as measures of the intra-individual variability of sleep. Of the 81 subjects included, 80 had seven nights of actigraphy data and one had six nights.

#### Self-Report Measures

Questionnaire-based, retrospective measures of sleep health were obtained at visit 1 using a selection of validated and commonly used inventories. The Pittsburgh Sleep Quality Index (PSQI) (Buysse et al. 1991) was used to assess sleep quality (global score) and self-reported habitual sleep duration (item 4), the Insomnia Severity Index (ISI) (Morin 1993) was used to assess insomnia symptoms, the Epworth Sleepiness Scale (ESS) was used to assess daytime sleepiness (Johns 1991), and the Horne-Östberg Morningness / Eveningness Questionnaire (MEQ) (Horne and Ostberg 1976) was used to assess chronotype preference. Participants were also asked to report their usual duration of sleep on weekdays (WD) versus weekends (WE), and the difference score was used to estimate self-reported sleep variability (WE-WD difference). Finally, the Chalder Fatigue Scale (CFS) was used to assess problems with fatigue in daily life (Chalder et al. 1993). For the PSQI, the global score was calculated where higher scores indicate poorer sleep quality. For the ISI and ESS, the sum score of all items was used, where higher scores indicate more problems with insomnia symptoms and daytime sleepiness. For the MEQ, the global score was calculated, where higher scores indicate a stronger preference for morningness. For alignment with actigraphy midpoint data, the MEQ global score was inverted in all correlation analyses such that higher scores indicate a preference for eveningness (later chronotype preference). For the CFS, the mean of all items was calculated, where a mean of 1 indicates no current problems with fatigue and higher scores indicate more problems.

#### Standardized Measure of Cognitive Control Function

To obtain a standardized measure of cognitive control function, participants completed the widely used Conners Continuous Performance Test 3 (CCPT-3) at visit 1 (Conners 2014). In this computerized task, a series of letters (A-Z) appear on the screen in random order, and participants are asked to press the spacebar everytime they see a letter, except for the letter X (“Not-X-CPT”). Participants were told to respond as quickly and accurately as possible. Using norms from the test provider (MHS Scoring Software, version 5.6.0, Multi-Health Systems Inc., Canada), T-scores for hit reaction time (hit RT), hit reaction time standard deviation (hit RT SD), commission errors (non-targets responded to), omission errors (targets missed), as well as the derivative measure detectability (d’), were extracted for a descriptive characterization of the participants.

### Neuroimaging Protocol and Data Acquisition

#### fMRI Task

To assess the neural correlates of cognitive control processing, we used a well-validated Not-X-CPT task which was specifically developed for a mixed block/event-related fMRI design and allows for the study of proactive and reactive cognitive control processing, including time-on-task effects (Olsen et al. 2013, 2015, 2018). Briefly, letters from A-Z were presented for 250 ms each, with an interstimulus interval varying between 1, 2 and 4 seconds. Participants were asked to respond whenever they saw a letter appear (targets), but to withhold their response when the letter X appeared (non-targets) (Figure 1b). They were told to respond as quickly and accurately as possible. Response speed and accuracy were recorded using fiber-optic response grips (Nordic Neurolabs, Bergen, Norway) held in the participants’ dominant hand (determined using the Edinburgh Handedness Inventory (Oldfield 1971)). In total, 480 stimuli were presented, and non-target (X) frequency was 10%. The task paradigm consisted of two task runs (duration ~15 minutes each), which were counterbalanced with regard to stimulus presentation to allow for investigation of time-on-task effects. The paradigm and stimulus presentation are described in greater detail elsewhere (Olsen et al. 2013). The task was presented on an HDMI monitor (Nordic Neurolabs, Bergen, Norway) via the EPrime 3.0 software (Psychology Software Tools, Pittsburgh, PA). Participants viewed the HDMI screen via a coil mounted mirror (Siemens, Erlangen, Germany). The following performance measures were extracted from the behavioral logs and used in further analyses: hit RT and hit RT SD in milliseconds (target stimuli), omission errors (targets missed), and commission errors (non-targets responded to).

#### Mental Fatigue and Sleepiness During fMRI Task Performance

To assess subjective levels of mental fatigue and sleepiness during fMRI task performance, participants were asked the following questions via the MRI speaker system halfway through the task: 1) “On a scale from 1 to 10, how mentally fatigued do you feel right now, where 1 equals ‘not at all’ and 10 equals ‘severely’?”, and 2) “On a scale from 1 to 10, how sleepy do you feel right now, where 1 equals ‘extremely alert’ and 10 equals ‘can’t keep awake’?”. The question about sleepiness was adapted from the 10-point version of the Karolinska Sleepiness Scale (KSS) (Akerstedt and Gillberg 1990; Shahid et al. 2011).

#### MRI Data Acquisition and Preprocessing

MRI data were acquired on a 3T Skyra scanner using a 32-channel head-coil (Siemens, Erlangen, Germany). For the task fMRI, two series of multiband T2*-weighted echo planar images (EPI) with whole-brain coverage were acquired (947 volumes; phase encoding direction = A-P; SMS = 6; TR = 0.970 s; TE = 34.2 ms; FA = 60°; voxel size = 2.5 mm isotropic, FOV = 260 mm). For correction of susceptibility induced distortions, two series of spin-echo planar images in opposite phase encoding directions (AP-PA) were acquired after each fMRI run (3 volumes; TR = 7.33 s; TE = 60.8 ms; FA = 90°; voxel size = 2.5 mm isotropic, FOV = 260 mm). For anatomical co-registration, a high-resolution, 3D T1-weighted MPRAGE volume was acquired (TR = 2.3 s; TE = 29.2 ms; FA = 9°, voxel size = 1 × 1 × 1.2 mm; FOV = 256 mm).

Anatomical T1-weighted (T1w) images were preprocessed using fMRIPrep version 20.2.3 (Esteban et al. 2018, 2019) (RRID:SCR_016216), which is based on Nipype 1.6.1 (Gorgolewski et al. 2011, 2018) (RRID:SCR_002502). The T1w images were corrected for intensity non-uniformity with N4BiasFieldCorrection (Tustison et al. 2010), distributed with ANTs 2.3.3 (Avants et al. 2008) (RRID:SCR_004757) and thereafter skull-stripped with a Nipype implementation of the antsBrainExtraction.sh workflow (from ANTs), using OASIS30ANTs as a target template. Brain tissue segmentation of cerebrospinal fluid (CSF), white-matter (WM) and gray-matter (GM) was performed on the brain-extracted T1w image using FAST (FSL 5.0.9, RRID:SCR_002823) (Zhang et al. 2001). Brain surfaces were reconstructed using recon-all (Dale et al. 1999) (FreeSurfer 6.0.1, RRID:SCR_001847), and the T1 brain mask was refined with a custom variation of the method to reconcile ANTs-derived and FreeSurfer-derived segmentations of the cortical gray-matter of Mindboggle (Klein et al. 2017) (RRID:SCR_002438). The resulting brain mask was used to run brain extraction on the T1w image using BET (Smith 2002), after mean dilation of non-zero voxels to ensure full brain coverage in the resulting brain-extracted image.

FMRI data pre-processing was carried out using FEAT (FMRI Expert Analysis Tool) Version 6.00, part of FSL (FMRIB’s Software Library, Oxford, UK) (Jenkinson et al. 2012). First, to improve registrations from functional to structural space, single-band reference images from the multiband EPI sequences were used, and susceptibility distortion correction was applied by running TOPUP (Andersson et al. 2003) on the spin-echo planar images, followed by b0 unwarping as implemented in FEAT. Registration of functional images to the preprocessed T1w image/the 2-mm MNI standard space template was carried out using FLIRT with boundary-based registration/12 degrees of freedom (Jenkinson and Smith 2001; Jenkinson et al. 2002) and further refined using FNIRT nonlinear registration with a 10-mm warp resolution (Andersson et al. 2007). The following pre-statistical processing steps were applied: motion correction using MCFLIRT (Jenkinson et al. 2002); non-brain removal using BET (Smith 2002); spatial smoothing using a Gaussian kernel of FWHM 6 mm and grand-mean intensity normalization of the entire 4D dataset by a single multiplicative factor. High-pass temporal filtering (Gaussian-weighted least-squares straight line fitting) was applied using sigmas of 50.0 s for block-related analyses, and 25.0 s for event-related analyses.

### Statistical Analysis

IBM SPSS Statistics version 27 was used to calculate the daily sleep midpoints, as well as 7-day averages and standard deviations for all actigraphy outcome measures. Actigraphy, fMRI performance and questionnaire data were then further analyzed and visualized in RStudio (version 1.4.1103; R version 4.0.3) using the *tidyverse* (Wickham et al. 2019), *lubridate* (Grolemund and Wickham 2011), *psycho* (Makowski 2018), *psych* (Revelle 2022) and *corrplot* (Wei and Simko 2021) packages. Functional MRI data were analyzed using FSL FEAT (FMRI Expert Analysis Tool) Version 6.00 (FMRIB’s Software Library, Oxford, UK).

#### fMRI Task Performance

Individual mean hit RT, hit RT SD, number of omissions, commissions, and detectability were calculated for use in further analyses. Detectability was calculated using the *psycho* package in R (Makowski 2018). Both overall task performance (collapsed across the two task runs) and time-on-task changes (Δ) were calculated for all measures. To explore associations between fMRI performance and the different measures of sleep health, a partial correlation analysis (adjusted for age, sex, and years of education, determined a-priori) was performed. Spearman’s correlation coefficient was used to account for non-normally distributed data. Given the explorative nature of this analysis, no formal correction for multiple comparisons was performed, and results should therefore be considered preliminary.

#### Whole-Brain fMRI Analyses

Prior to statistical analysis, all individual task fMRI runs were checked for excessive motion. The values for relative root mean square displacement were very low (mean: 0.1 mm: max: 0.25 mm), and all subjects were therefore included in subsequent analyses. Single-subject general-linear models (GLM) were first applied for each task run (run 1 and run 2), using FSL’s FILM with local autocorrelation correction (Woolrich et al. 2001). The following contrasts were computed for use in the primary analysis (1): task blocks > fixation blocks (Proactive Cognitive Control), and non-targets (X) > targets (A-Z) (Reactive Cognitive Control). Contrasts from individual task runs were then combined for each participant using a fixed effects model, to compute mean activation across runs. For the secondary analysis (2), to investigate time-on-task effects, each task run was divided into four time epochs in which Proactive and Reactive Cognitive Control BOLD activations were estimated. Mean time-on-task change across runs (Δ Proactive Cognitive Control and Δ Reactive Cognitive Control) was then computed for each participant using a fixed effects model, by contrasting time epoch 1 with time epoch 4.

Linear whole-brain associations between lower-level contrast estimates and sleep health measures were modeled using a mixed-effects model (FLAME 1 + 2) (Woolrich et al. 2004). Separate GLMs were run to test (1) associations between Proactive- and Reactive Cognitive Control and sleep health, and (2) Δ Proactive- and Δ Reactive Cognitive Control and sleep health. The following sleep health measures were demeaned and included as covariates of interest in separate models (one per fMRI contrast): actigraphy-derived mean sleep duration, sleep duration SD, mean midpoint, midpoint SD, sleep efficiency, and SOL; self-reported sleep duration (PSQI), chronotype (MEQ inverse global score), sleep variability (WE-WD difference), sleep quality (PSQI), insomnia symptoms (ISI), problems with fatigue in daily life (CFS), daytime sleepiness (ESS), mental fatigue during the task, and sleepiness during the task (KSS). Additionally, (3) group-level average activation for all BOLD contrasts (excluding sleep health covariates) were modeled in a supplementary analysis, to provide context for primary/secondary results, and for evaluation of the validity and replicability of the fMRI protocol.

All models were adjusted for age, sex, years of education, and fMRI task performance (hit RT, omissions and commissions), to control for confounding factors related to sleep health, cognitive control function, and BOLD signal (Olsen et al. 2015; Shanmugan and Satterthwaite 2016; Gurvich et al. 2018; Evans et al. 2021; van de Langenberg et al. 2022). Adjustment for fMRI task performance was applied as we were mainly interested in differences in neuronal processing during cognitive control related to sleep health, as opposed to differences in BOLD activation caused by mere behavioral variability - which may be related to a range of personal and/or contextual factors (Price et al. 2006; Yarkoni et al. 2009; Grinband et al. 2011). Models of Proactive and Reactive Cognitive Control were adjusted for overall (whole-task) performance, and models of Δ Proactive Cognitive Control and Δ Reactive Cognitive Control (time-on-task change) were adjusted using performance Δ scores. The six head motion parameters from FEAT were also added to each model as separate regressors.

To control the family-wise error (FWE) rate, in each individual model, cluster-based inference based on Gaussian Random Field Theory (RFT) was applied using a cluster-defining threshold of Z > 3.1, and a cluster probability threshold of p < .05 (Worsley 2011). For statistically significant results, the size (number of voxels), P-value, maximum Z-values, and coordinates in standard 2×2×2 MNI space for significant clusters were extracted for descriptive purposes. The minimum significant cluster size, i.e., the minimum number of contiguous voxels (Z > 3.1) required for a cluster to be considered significant (p < .05), is also reported for each model. Anatomical locations were determined using the FSLeyes software, version 1.3.0, with the incorporated Harvard Oxford cortical and subcortical structural brain atlases and visual inspection.

## Results

### Demographics and Standardized Cognitive Control Function

An overview of demographic variables and scores on the standardized assessment of cognitive control function (CCPT-3) can be found in Table 1. For the CCPT-3, lower scores reflect better performance. On the group level, participants had relatively faster reaction times while maintaining an expected number of errors as compared to the norm group (hit RT: mean T-score = 43.01, SD = 6.09; commission errors: mean T-score = 50.99, SD = 9.53).

**Table 1.** Demographics and Standardized Cognitive Control Function

| Variable | n | Mean (SD) | Min | Max |
| --- | --- | --- | --- | --- |
| **Demographics** |  |  |  |  |
| Female / Male | 41 / 40 | n/a | n/a | n/a |
| Age (years) | 81 | 27.82 (5.42) | 20.96 | 39.58 |
| Years of education<sup>a</sup> | 81 | 16.28 (2.39) | 12 | 21 |
| **Standardized Cognitive Control Function** |  |  |  |  |
| **(Conners CPT-3, T-scores)** |  |  |  |  |
| Hit RT, mean | 81 | 43.01 (6.09) | 33 | 63 |
| Hit RT, SD | 81 | 41.49 (5.84) | 32 | 66 |
| Omission errors | 81 | 45.60 (2.72) | 43 | 60 |
| Commission errors | 81 | 50.99 (9.53) | 36 | 78 |
| Detectability (d') | 81 | 47.49 (7.39) | 32 | 66 |

The standardized scores on the CPT-3 have a mean of 50 and a standard deviation of 10. Higher scores indicate poorer performance, and lower scores indicate better performance. <sup>a</sup>Years of education = the number of years corresponding to the highest completed level *or* ongoing level of education (educational attainment). SD = standard deviation; RT = reaction time.

### Measures of Sleep Health

Table 2 provides an overview of measures of sleep health. The mean objective sleep duration (averaged over 7 days) was 7.21 hours (SD = 0.66 hours), which is within the range of recommended sleep duration for adults (7 - 8 hours) (Hirshkowitz et al. 2015). The mean sleep efficiency and SOL also indicated overall good quality sleep (≥85% and <30 minutes, respectively) (Ohayon et al. 2017). For sleep timing, the mean midpoint was at 03:52 AM, and mean of scores on the MEQ was 54.29 (SD = 8.86). The distribution of scores on the MEQ showed that a majority of participants fell within the ‘intermediate’ type, with a slight lean toward ‘moderately morning’ (Horne and Ostberg 1976). Finally, the mean scores on the PSQI and ISI suggest a low prevalence of sleep problems (using a cut-off for ‘normal sleep’ of 5 for the PSQI, and 7 for the ISI (Buysse et al. 2008; Morin et al. 2011)). Taken together, the results show that participants had good sleep overall, but that there was inter-individual variability as expected within the normal range.

**Table 2.**
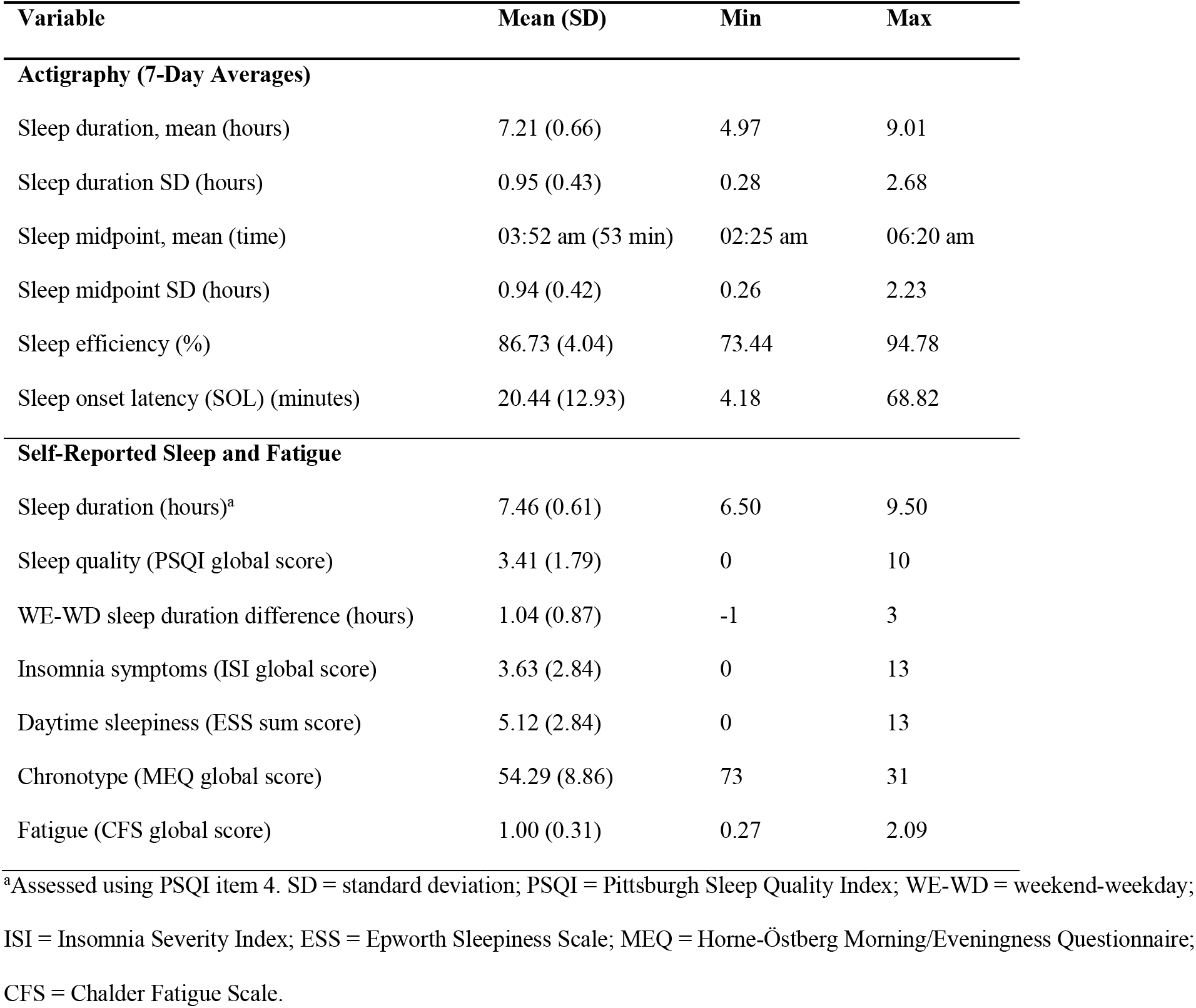
Sleep Health Measures (N = 81)

| Variable | Mean (SD) | Min | Max |
| --- | --- | --- | --- |
| **Actigraphy (7-Day Averages)** |  |  |  |
| Sleep duration, mean (hours) | 7.21 (0.66) | 4.97 | 9.01 |
| Sleep duration SD (hours) | 0.95 (0.43) | 0.28 | 2.68 |
| Sleep midpoint, mean (time) | 03:52 am (53 min) | 02:25 am | 06:20 am |
| Sleep midpoint SD (hours) | 0.94 (0.42) | 0.26 | 2.23 |
| Sleep efficiency (%) | 86.73 (4.04) | 73.44 | 94.78 |
| Sleep onset latency (SOL) (minutes) | 20.44 (12.93) | 4.18 | 68.82 |
| **Self-Reported Sleep and Fatigue** |  |  |  |
| Sleep duration (hours)<sup>a</sup> | 7.46 (0.61) | 6.50 | 9.50 |
| Sleep quality (PSQI global score) | 3.41 (1.79) | 0 | 10 |
| WE-WD sleep duration difference (hours) | 1.04 (0.87) | -1 | 3 |
| Insomnia symptoms (ISI global score) | 3.63 (2.84) | 0 | 13 |
| Daytime sleepiness (ESS sum score) | 5.12 (2.84) | 0 | 13 |
| Chronotype (MEQ global score) | 54.29 (8.86) | 73 | 31 |
| Fatigue (CFS global score) | 1.00 (0.31) | 0.27 | 2.09 |

<sup>a</sup>Assessed using PSQI item 4. SD = standard deviation; PSQI = Pittsburgh Sleep Quality Index; WE-WD = weekend-weekday; ISI = Insomnia Severity Index; ESS = Epworth Sleepiness Scale; MEQ = Horne-Östberg Morning/Eveningness Questionnaire; CFS = Chalder Fatigue Scale.

### fMRI Task Performance and Associations With Sleep Health

Behavioral data from the fMRI task are presented in Figure 3 and Supplementary Table 1. Data are presented both for the task as a whole and for time-on-task changes (Δ).

**Figure 3.**
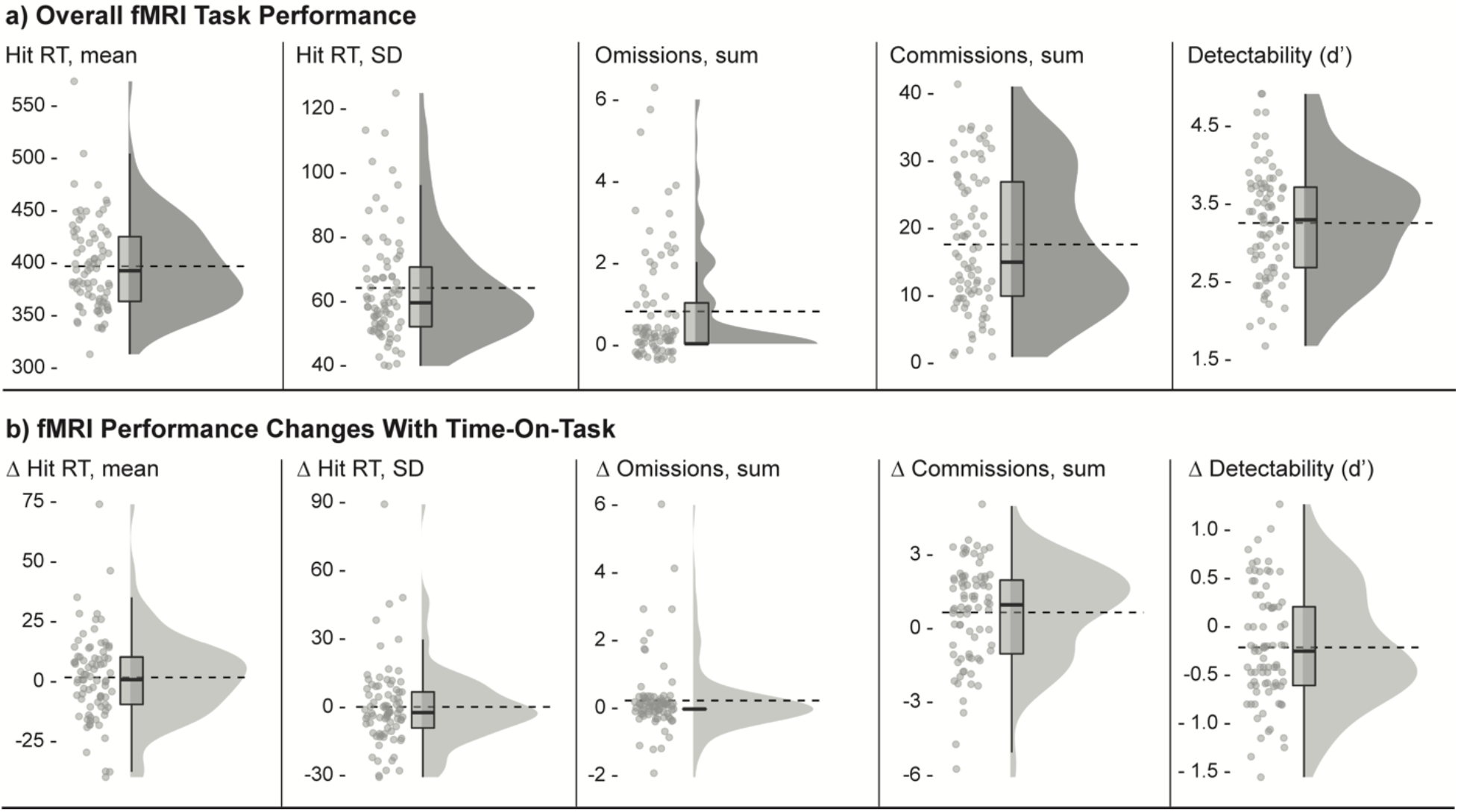
fMRI Task Performance. Individual means for each outcome variable are plotted as raincloud plots (Allen et al. 2021) with overlaid boxplots, as well as dashed horizontal lines indicating group means. Figure a) shows overall task performance and b) shows performance change scores with time on task (time epoch 4 - time epoch 1). Hit RT and hit RT SD refer to target responses (letters A - Z). Omissions refer to missed target letters (A-Z), and commissions refer to pushed non-targets (X). Detectability (d’) refers to the ability to discriminate targets from non-targets. fMRI = functional magnetic resonance imaging, RT = reaction time, SD = standard deviation.

Results from an explorative investigation of associations between fMRI task performance and sleep health variables are presented in Figure 4. Given the explorative nature of this analysis, no formal correction for multiple comparisons was performed, and the results should therefore be considered preliminary. Significant correlations (at *α* = .05, uncorrected) are labeled according to their p-value in the figure (* = p < .05, ** = p < .01, *** = p < .001), and summarized here. Most associations were observed for actigraphy-derived sleep duration: shorter habitual duration was associated with poorer task performance overall (higher hit RT SD, more errors, and lower detectability) as well as with increased time on task (longer hit RT, higher hit RT SD, and lower detectability). More variable habitual sleep duration (higher SD) was also associated with more commission errors and lower detectability. Further, shorter *self-reported* sleep duration was associated with poorer performance with time on task (more omission errors and lower detectability), relatively later chronotype preference was associated with lower hit RT SD with time on task, and higher levels of daytime sleepiness were associated with more omissions and longer hit RTs with time on task. There were no other statistically significant associations (p < 0.05), between fMRI task performance and sleep health measures.

**Figure 4.**
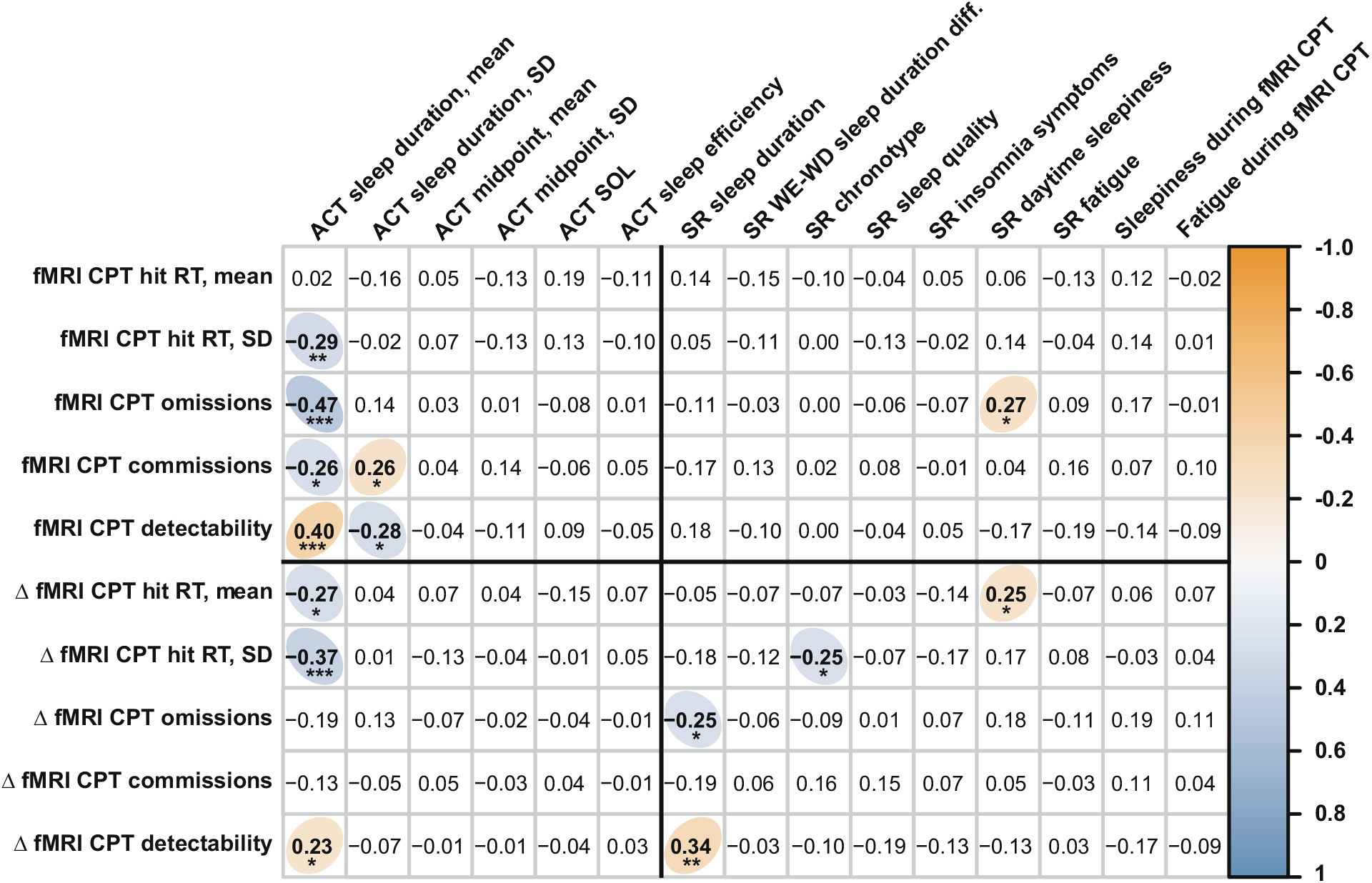
Partial Correlations between Task fMRI Performance and Sleep Health Measures. The correlogram depicts partial correlation coefficients (Spearman’s rho; adjusted for age, sex, and education) between fMRI task performance measures (Y axis) and the different sleep health measures (X axis). Measures of overall task performance are listed above the horizontal black line and measures of time-on-task changes (Δ) are listed below the line. Objective measures of sleep (actigraphy-derived) are listed to the left of the vertical black line and self-report measures are listed to the right of the line. Statistically significant correlations (p < .05, not corrected for multiple comparisons) are marked with colored ellipses (positive correlations in orange and negative correlations in blue). Given the explorative purpose of this analysis, no formal correction for multiple comparisons was performed, and results should therefore be considered preliminary. To indicate which findings would survive stricter statistical thresholds, significant correlations have been labeled according to their uncorrected p-value (* = p < .05, ** = p < .01, *** = p < .001). ACT = actigraphy; SR = self-reported; fMRI = functional magnetic resonance imaging; CPT = continuous performance test; RT = reaction time, SD = standard deviation.

### Whole-Brain fMRI Analyses

#### Associations Between Proactive and Reactive Cognitive Control Activations and Sleep Health

Statistically significant associations between cognitive control activations and sleep health measures are presented in Figures 5, 6 and in Table 3. Most were found for Reactive Cognitive Control activation.

**Figure 5.**
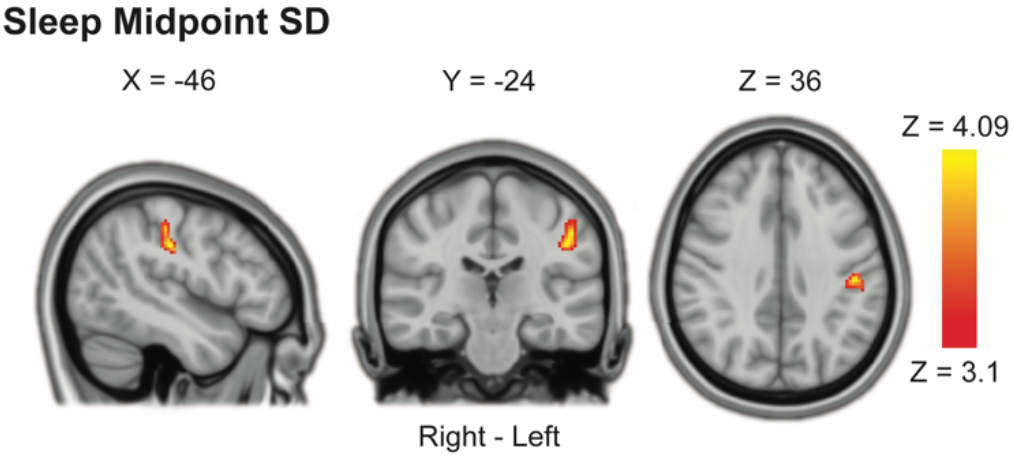
Association between Proactive Cognitive Control Processing and Sleep Midpoint SD. More variable sleep midpoint (midpoint SD) was associated with stronger proactive cognitive control activation in the left postcentral gyrus. Results were obtained using mixed-effects models and are presented on a 1-mm MNI standard space template. Cluster-based inference was used to control the family-wise error rate in each model (cluster-defining threshold = Z > 3.1, cluster probability threshold = p < .05). Slices that best represent the cluster have been selected. As these are 2D representations of 3D volumes, the cluster may only be partly visible (see Table 3 for details on cluster size/coordinates). SD = standard deviation, MNI = Montreal Neurological Institute.

**Figure 6.**
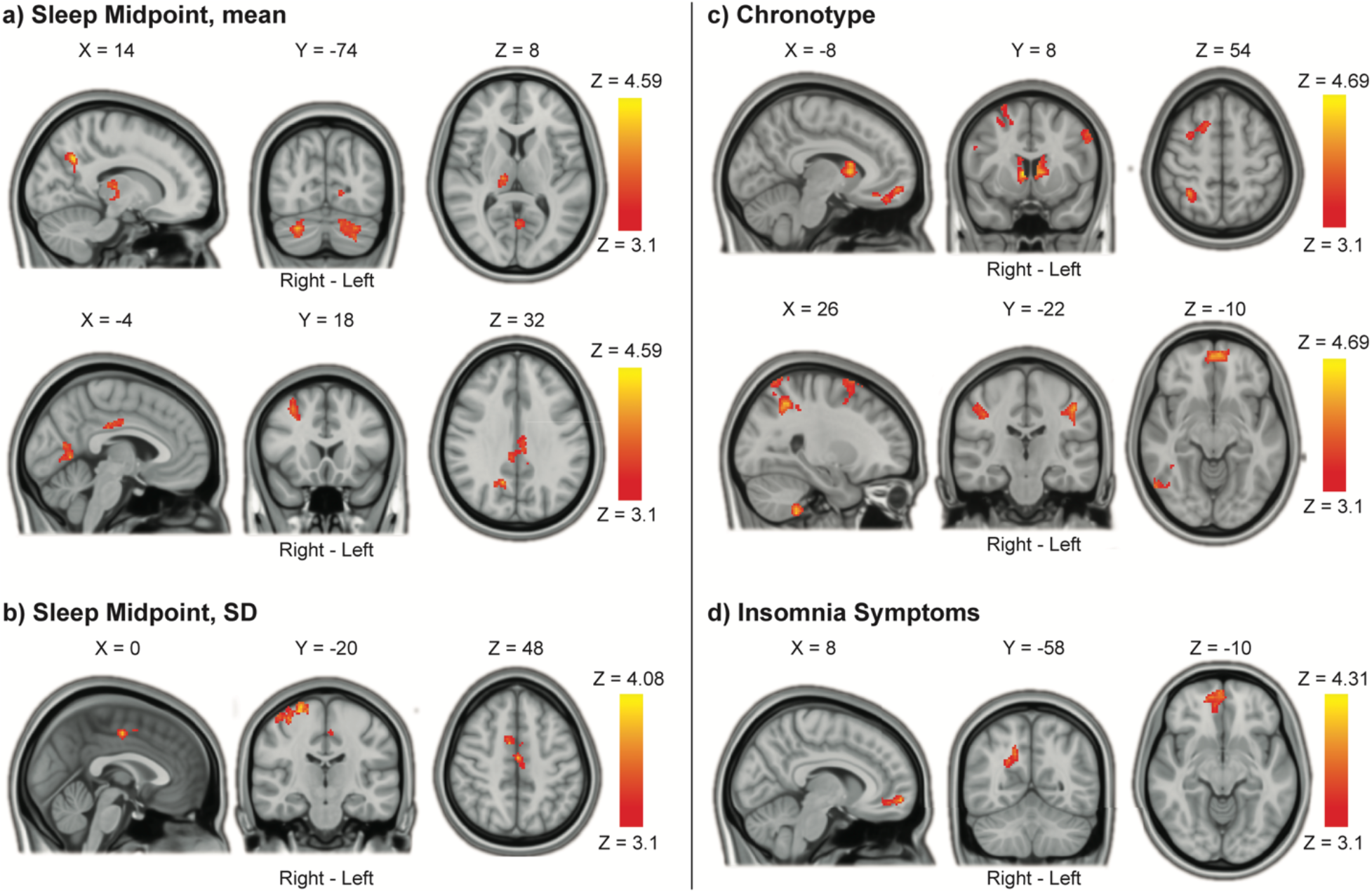
Associations between Reactive Cognitive Control Processing and Sleep Health. a) Later sleep midpoint was associated with higher reactive cognitive control activations in the cerebellum, lingual gyrus, thalamus, right precuneus cortex, posterior cingulate gyrus, and right middle frontal gyrus. b) More variable sleep midpoint (midpoint SD) was associated with higher reactive cognitive control activation in the precentral / postcentral gyrus, and in the juxtapositional lobule (supplementary motor area). c) Later chronotype preference was associated with widespread higher reactive cognitive control activation, including frontal, parietal and temporal cortex as well as the cerebellum, caudate and thalamus. d) Higher levels of insomnia symptoms were associated with higher reactive cognitive control processing in the right precuneus and ventromedial frontal cortex. Results were obtained using mixed-effects models and are presented on a 1-mm MNI standard space template. Cluster-based inference was used to control the family-wise error rate in each model (cluster-defining threshold = Z > 3.1, cluster probability threshold = p < .05). Slices that are most representative for the overall findings (anatomical, and across different clusters) have been selected. As these are 2D representations of 3D volumes, some of the clusters may only be partly visible (see Table 3 for details on cluster size/coordinates). SD = standard deviation, MNI = Montreal Neurological Institute.

**Table 3.** Associations Between Cognitive Control Activations and Sleep Health

| Contrast / Region | Right | Size | P Value | Peak Z | Peak Coordinates |  |  |
| --- | --- | --- | --- | --- | --- | --- | --- |
|  | / Left | (#voxels) |  | Value | (MNI) |  |  |
|  |  |  |  |  | X | Y | Z |
| Proactive Cognitive Control: | Minimum significant cluster size: 182 voxels |  |  |  |  |  |  |
| Higher Sleep Midpoint SD |  |  |  |  |  |  |  |
| Postcentral Gyrus | L | 198 | .0366 | 4.09 | -46 | -24 | 36 |
| Reactive Cognitive Control: | Minimum significant cluster size: 168 voxels |  |  |  |  |  |  |
| Later Sleep Midpoint |  |  |  |  |  |  |  |
| Cerebellum: Right Crus I | R | 332 | .0020 | 4.22 | 28 | -74 | -32 |
| Cerebellum: Left Crus I & II | L | 306 | .0032 | 4.08 | -18 | -70 | -36 |
| Cingulate Gyrus, posterior division | R | 266 | .0068 | 4.37 | 4 | -36 | 26 |
| Lingual Gyrus / Intracalcarine Cortex | L | 248 | .0096 | 4.37 | -6 | -62 | 2 |
| Precuneus Cortex | R | 235 | .0123 | 4.38 | 14 | -60 | 36 |
| Thalamus | R | 226 | .0148 | 3.98 | 14 | -20 | 8 |
| Middle Frontal Gyrus | R | 210 | .0204 | 4.59 | 32 | 12 | 52 |
| Reactive Cognitive Control: | Minimum significant cluster size: 168 voxels |  |  |  |  |  |  |
| Higher Sleep Midpoint SD |  |  |  |  |  |  |  |
| Precentral & Postcentral Gyri | R | 592 | < .0001 | 4.08 | 24 | -18 | 74 |
| Juxtapositional Lobule | R | 169 | .0488 | 3.85 | 0 | -12 | 48 |
| Reactive Cognitive Control: | Minimum significant cluster size: 167 voxels |  |  |  |  |  |  |
| Later Chronotype Preference |  |  |  |  |  |  |  |
| Superior Parietal Lobule | R | 760 | < .0001 | -4.69 | 38 | -58 | 68 |
| Cerebellum: Right VIIIb | R | 709 | < .0001 | -4.40 | 26 | -42 | -54 |
| Caudate | R / L | 495 | .0001 | -4.68 | 8 | 8 | 4 |
| Middle Temporal Gyrus, temporooccipital part | R | 367 | .0010 | -4.38 | 48 | -52 | 2 |
| Superior Frontal Gyrus | R | 311 | .0028 | -4.41 | 24 | 14 | 72 |
| Inferior Frontal Gyrus, pars opercularis | R | 282 | .0047 | -4.03 | 40 | 10 | 22 |
| Frontal Pole / Frontal Medial Cortex | R / L | 273 | .0056 | -4.22 | -4 | 58 | -10 |
| Postcentral Gyrus | L | 252 | .0085 | -4.38 | -46 | -18 | 30 |
| Postcentral Gyrus | R | 242 | .0103 | -3.95 | 40 | -24 | 38 |
| Precentral Gyrus | L | 228 | .0136 | -4.08 | -52 | 6 | 42 |

**Table 3 Continued**
| Contrast / Region | Right | Size | P Value | Peak Z | Peak Coordinates |  |  |
| --- | --- | --- | --- | --- | --- | --- | --- |
|  | / Left | (#voxels) |  | Value | (MNI) |  |  |
|  |  |  |  |  | X | Y | Z |
| Reactive Cognitive Control | Minimum significant cluster size = 168 voxels |  |  |  |  |  |  |
| More Insomnia Symptoms |  |  |  |  |  |  |  |
| Precuneus Cortex | R | 247 | .0097 | 3.81 | 16 | -58 | 36 |
| Frontal Pole / Frontal Medial Cortex | R | 201 | .0245 | 4.31 | 4 | 58 | -8 |

For Proactive Cognitive Control, a more variable sleep midpoint (higher midpoint SD) was associated with higher BOLD activation in the left postcentral gyrus (Figure 5). None of the other sleep health variables was associated with overall Proactive Cognitive Control activation.

For Reactive Cognitive Control, later sleep midpoint was associated with greater BOLD activation in the cerebellum, lingual gyrus, thalamus, right precuneus cortex, posterior cingulate gyrus, and right middle frontal gyrus (Figure 6a). A more variable sleep midpoint (higher midpoint SD) was associated with greater Reactive Cognitive Control in the right precentral / postcentral gyrus, as well as the juxtapositional lobule (supplementary motor area) (Figure 6b). Later chronotype preference was associated with greater Reactive Cognitive Control activation in widespread regions encompassing frontal, parietal, and temporal cortex as well as the cerebellum, caudate and thalamus (Figure 6c). Finally, a higher level of insomnia symptoms was associated with greater Reactive Cognitive Control activation in the right precuneus and ventromedial frontal cortex (Figure 6d).

#### Associations Between Δ Proactive Cognitive Control, Δ Reactive Cognitive Control and Sleep Health

Statistically significant associations between time-on-task changes in cognitive control activation and sleep health measures are presented in Figures 7, 8 and in Table 4. Most associations with time-on-task effects were found for Δ Proactive Cognitive Control activation.

**Figure 7.**
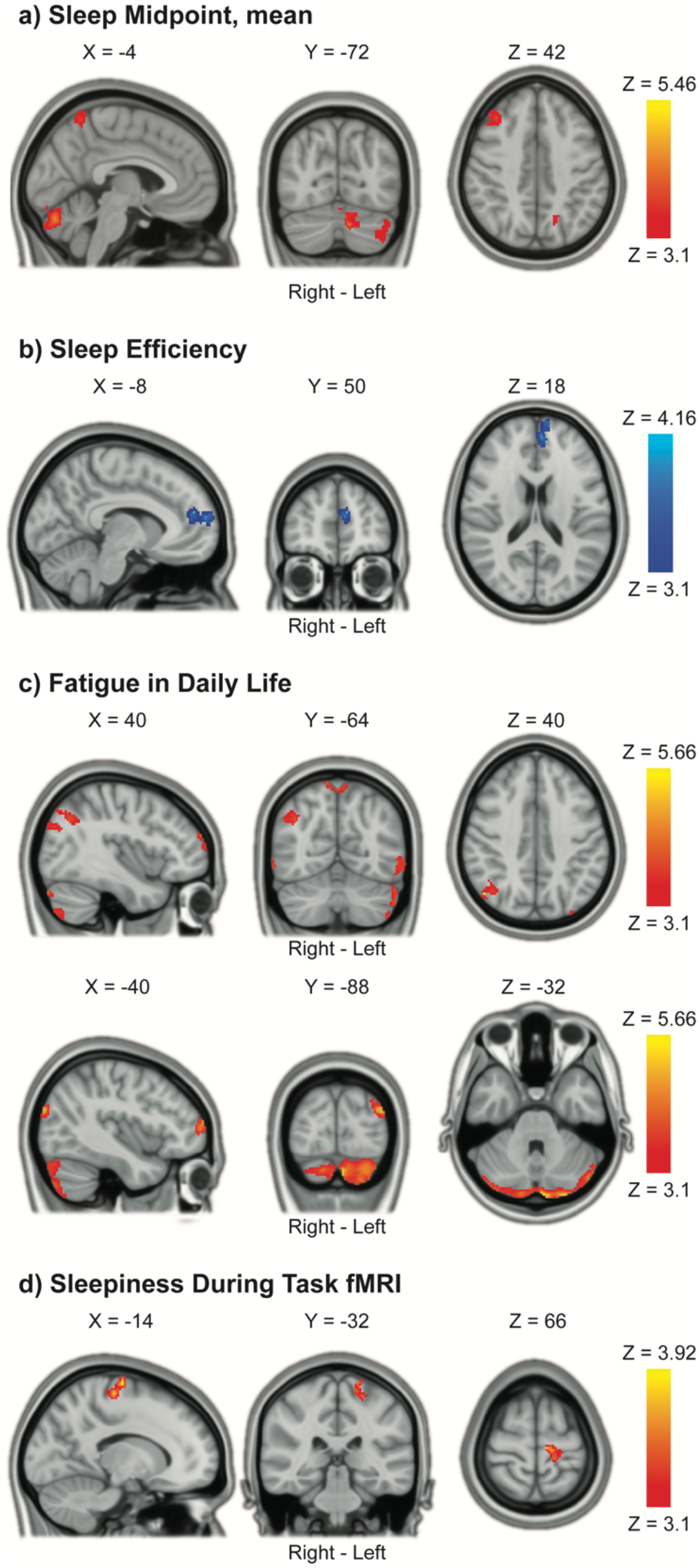
Time-On-Task (TOT) Increases in Proactive Cognitive Control Associated With Sleep Health. a) Later sleep midpoint was associated with increased Proactive Cognitive Control activations in the cerebellum, precuneus cortex and right middle frontal gyrus with TOT. b) Lower sleep efficiency was associated with increased Proactive Cognitive Control activations in the paracingulate gyrus / left frontal pole with TOT. c) More problems with fatigue in daily life were associated with increased Proactive Cognitive Control activations in widespread areas of the brain, including the cerebellum, occipital cortex, precuneus and frontal pole with TOT. d) Higher levels of sleepiness during fMRI task performance were associated with increased Proactive Cognitive Control activations in the left precentral / postcentral gyrus with TOT. Results were obtained using mixed-effects models and are presented on a 1-mm MNI standard space template. Cluster-based inference was used to control the family-wise error rate in each model (cluster-defining threshold = Z > 3.1, cluster probability threshold = p < .05). Slices that are most representative for the overall findings (anatomical, and across different clusters) have been selected. As these are 2D representations of 3D volumes, some of the clusters may only be partly visible (see Table 4 for details on cluster size/coordinates). TOT = time on task, SD = standard deviation; MNI = Montreal Neurological Institute.

**Figure 8.**
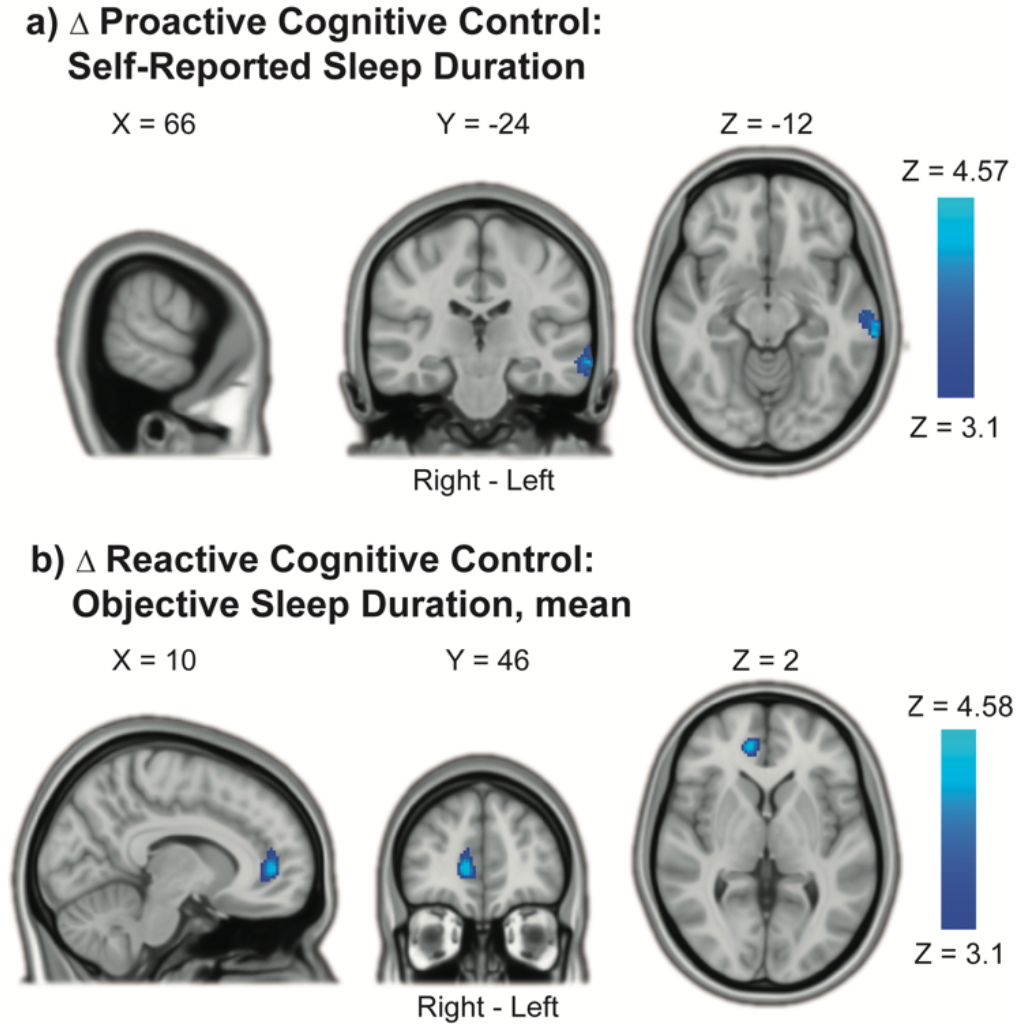
Time-On-Task (TOT) Decreases in Cognitive Control Processing with Shorter Sleep Duration. a) Shorter self-reported sleep duration was associated with decreased Proactive Cognitive Control activations in the left middle temporal gyrus with TOT. b) Shorter objective sleep duration (7-day mean) was associated with decreased Reactive Cognitive Control activation in the right paracingulate / anterior cingulate gyrus with TOT. Results were obtained using mixed-effects models and are presented on a 1-mm MNI standard space template. Cluster-based inference was used to control the family-wise error rate in each model (cluster-defining threshold = Z > 3.1, cluster probability threshold = p < .05). Slices that best represent the clusters have been selected. As these are 2D representations of 3D volumes, the clusters may only be partly visible (see Table 4 for details on cluster size/coordinates). TOT = time on task, MNI = Montreal Neurological Institute.

**Table 4.**
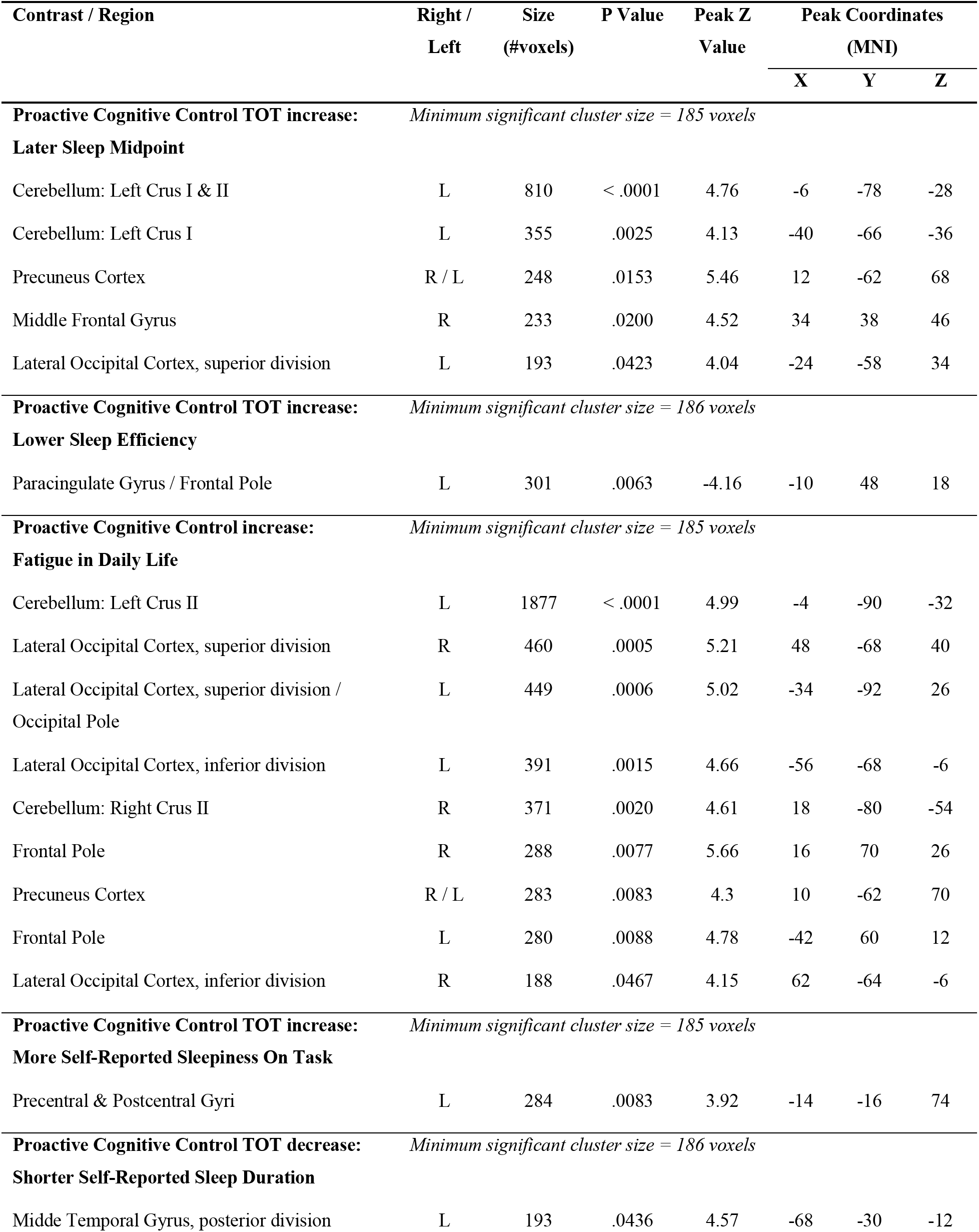
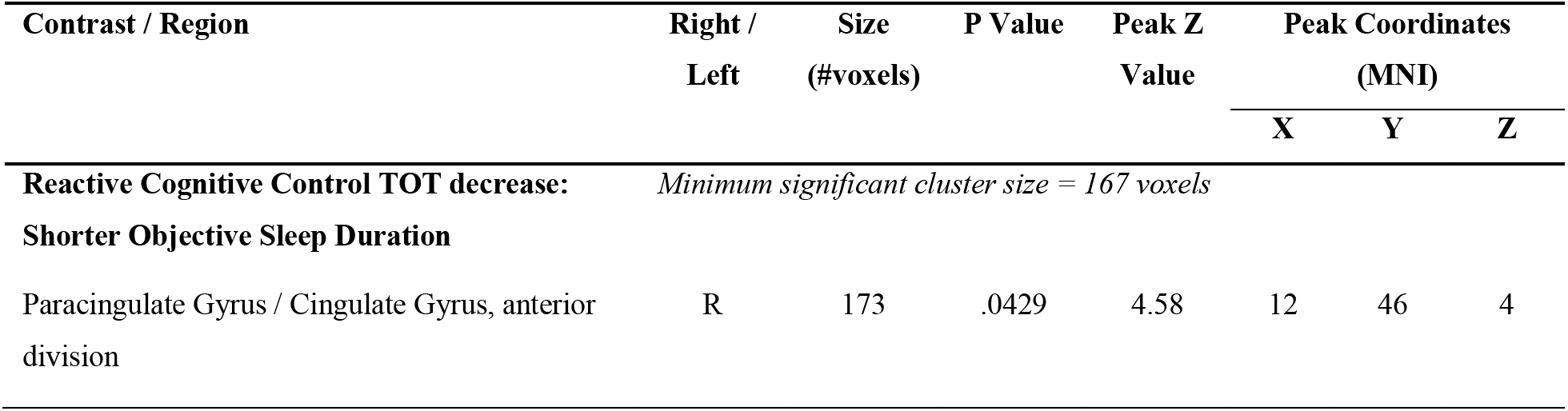
Associations Between Time-On-Task Change in Cognitive Control Activations and Sleep Health

For Δ Proactive Cognitive Control, later sleep midpoint was associated with increased BOLD activation with time on task in the cerebellum, precuneus cortex, and right middle frontal gyrus (Figure 7a). Lower sleep efficiency was associated with increased Proactive Cognitive Control activation in the left paracingulate gyrus / frontal pole with time on task (Figure 7b). More problems with fatigue in daily life were associated with increased Proactive Cognitive Control activation with time on task in widespread areas, including bilateral cerebellum, occipital cortex, precuneus cortex, and frontal pole (Figure 7c). More self-reported sleepiness during task performance was associated with increased Proactive Cognitive Control activation with time on task in the left precentral / postcentral gyrus (Figure 7d). Finally, shorter self-reported sleep duration was associated with *decreased* Proactive Cognitive Control activation with time on task in the left middle temporal gyrus (Figure 8a).

For Δ Reactive Cognitive Control, shorter objective (actigraphy-derived) sleep duration was associated with decreased BOLD activation with time on task in the right paracingulate / anterior cingulate gyrus (Figure 8b). None of the other sleep health measures was significantly associated with Δ Reactive Cognitive Control.

#### Supplementary fMRI Analysis: Group Average BOLD Activations

For transparency and evaluation of the validity (replicability) of the fMRI protocol, and to provide context for the discussion of results, we include group average activations for the contrasts used in our study (activation without sleep health covariates) in Supplementary Material (Figure 1, Tables 2, 3). Despite using a different scanner and fMRI sequence, group average activations were as expected and highly similar to previous studies (Olsen et al. 2013, 2015, 2018). The contrasts (1) for Proactive and Reactive Cognitive Control demonstrated robust and widespread BOLD activations, which converged on fronto-parietal regions, the insular cortex, dorsal striatum, thalamus, and the cerebellum - indicating core regions for cognitive control processing as observed in previous studies (Dosenbach et al. 2008; Niendam et al. 2012; Olsen et al. 2013). Contrasts for (2) time-on-task effects demonstrated a *decrease* in Proactive Cognitive Control activation within core control regions with increased time on task (which are typically ‘task-positive’ regions), and *increased* Proactive Cognitive Control activations in the precuneus, medial prefrontal cortex, and middle temporal gyrus (typically ‘task-negative’ regions). Meanwhile, there was *increased* Reactive Cognitive Control activation in core cognitive control areas with time on task, largely mirroring regions displaying *decreased* Proactive Cognitive Control activation. This group-level change in activity patterns suggests a relative shift from Proactive to Reactive Cognitive Control processing with increasing time on task.

## Discussion

In this prospective study of adult, normal sleepers, we observed multiple associations between cognitive control processing and habitual sleep health. Using whole-brain fMRI analyses, we found that measures indicating poorer sleep health were predominantly associated with stronger and more widespread BOLD activations during cognitive control processing (adjusted for age, sex, years of education and Not-X-CPT performance). Analyses focused on different temporal aspects of cognitive control processing yielded unique and complementary results. Later and more variable sleep midpoints, a relatively later chronotype preference, and higher levels of insomnia symptoms were associated with stronger *reactive* cognitive control activations. Furthermore, later sleep midpoints, lower sleep efficiency, more problems with fatigue in daily life, and more sleepiness during task performance were associated with increased *proactive* cognitive control activations with increasing time on task, reflecting more neural recruitment with time. Taken together, our results show that cognitive control function is linked to different aspects of habitual sleep health even in high-functioning, normal sleepers.

Given that fMRI task performance was adjusted for in our analyses (Price et al. 2006; Yarkoni et al. 2009), the dominating pattern of increased BOLD activations indicates that individuals with relatively poorer sleep health need to recruit more neuronal resources to support cognitive control function, which may reflect compensatory mechanisms and / or less efficient neural processing (Drummond et al. 2005; Chee and Tan 2010; Schmidt et al. 2015; Maire et al. 2018; Olsen et al. 2020). Shorter sleep duration was associated with *decreased* cognitive control activations with time on task as well as poorer task performance, suggesting an exaggerated, negative time-on-task effect in habitually shorter sleepers - perhaps reflecting a lower ability to maintain cognitive control function over longer periods of time. This fits well with previous evidence demonstrating that experimentally induced sleep loss exacerbates general time-on-task effects, and that the two share common neural substrates (Asplund and Chee 2013; Satterfield et al. 2017; Hudson et al. 2020).

Later and more variable sleep timing, later chronotype preference, and more insomnia symptoms were associated with stronger *reactive* cognitive control activations. This indicates that poorer sleep health is predominantly associated with a ‘hyper-reactive’ brain state, possibly due to increased recruitment of cognitive control resources as a response to conflicting (non-target) stimuli. This may indicate increased recruitment of cognitive control resources as a response to conflicting (non-target) stimuli. Increased reactive cognitive control has previously been associated with higher levels of stress (Husa et al. 2022) and anxiety (Fales et al. 2008; Schmid et al. 2015; Yang et al. 2018) in healthy individuals, and with poorer white matter organization, lower fluid intelligence, and more anxiety problems in adults born preterm (Olsen et al. 2018). The reactive hyperactivations observed here therefore suggest sub-optimal cognitive control processing in those with relatively poorer sleep health, and may point to a potential mechanism linking poorer sleep health to poorer mental health outcomes.

Beyond stronger activations in core cognitive control regions, later chronotype preference and more insomnia symptoms were also associated with higher activations in the ventromedial prefrontal cortex. The ventromedial prefrontal cortex has previously been implicated in self-referential cognition (Jenkins and Mitchell 2011; Abraham 2013) and processing of predictive value and reward (O’Doherty 2004; Etkin et al. 2011). Heightened activations here may therefore indicate experiencing the conflicting stimuli (non-targets) as more salient, and/or being more self-aware during conflict processing. The association with insomnia symptoms is also interesting in light of the ‘hyperarousal’ hypothesis, which postulates that insomnia disorder is linked to higher interoceptive awareness (Wei and Van Someren 2020), paired with an increased tendency to ruminate about sleep problems (Riemann et al. 2010; Fasiello et al. 2022). It is important to note that our findings are linked to insomnia *symptoms* in healthy participants, and not to the clinical diagnosis of insomnia disorder. However, the observed patterns of altered brain activity may still shed light on neural substrates underlying the development and maintenance of sleep problems (Nicolazzo et al. 2021; Faaland et al. 2022).

Analyses of time-on-task effects showed that measures of poorer sleep health were predominantly associated with increased *proactive* cognitive control activations in fronto-parietal and cerebellar regions. Mean group-level activation indicated a shift toward more *reactive* cognitive control processing with time on task, with decreasing proactive- and increasing reactive cognitive control activations in core control regions (Supplementary Figure 1c, 1d, 1e). Proactive cognitive control processing is believed to be more resource-demanding than reactive cognitive control (having a higher metabolic cost) (Burgess and Braver 2010; Braver 2012). The group-level shift toward more reactive cognitive control processing with time on task may therefore reflect an adaptive or more cost-efficient response - freeing up neural resources (lowering metabolic cost) and allowing participants to engage in other thought processes while still maintaining a satisfactory performance (Petersen et al. 1998; Neubauer and Fink 2009). Hence, the time-on-task increases in proactive cognitive control activations observed with poorer sleep health may indicate compensatory mechanisms, i.e., recruitment of relatively more neural resources in order to maintain cognitive control function over time (Olsen et al. 2015, 2020).

The strongest association for time-on-task effects was linked to problems with fatigue in daily life, for which there were widespread BOLD increases in cerebellar, occipital, and prefrontal areas with longer time on task. Heightened task-related BOLD activity has previously been observed in patient groups suffering from chronic fatigue, as compared to healthy controls (Cook et al. 2007; DeLuca et al. 2009; Almutairi et al. 2020). Our findings therefore imply that analyses of time-on-task effects are sensitive to temporal changes in neural activity associated with subtle symptoms also in healthy individuals. We did not observe statistically significant associations between time-on-task changes and self-reported fatigue during task performance. There was, however, an association with self-reported sleepiness during the task. Fatigue research has repeatedly demonstrated a distinction between fatigue and sleepiness, as well as ‘trait’ (stable over time) versus ‘state’ (momentary) measures (Kluger et al. 2013; Wylie et al. 2022). Our findings mirror this phenomenon, showing a widespread increase in brain activity related to ‘trait’ fatigue (as measured by the CFS), no evidence for differences in brain activity related to ‘state’ fatigue (mental fatigue during task performance), and unique activity changes related to task-related sleepiness centered in motor areas. One possible interpretation of these findings is that, while the time-on-task increases in BOLD activity may be compensatory in the moment, they might come with a longer-term cost (higher levels of trait fatigue) (Kohl et al. 2009; Olsen et al. 2015).

Shorter sleep duration was associated with *decreased* cognitive control activations with time-on-task (Figure 8). This finding is in contrast to the general pattern of increased activations with poorer sleep health in our study, but aligns well with existing evidence linking shorter sleep and sleep loss with lower task-related BOLD activations (Krause et al. 2017; Uy and Galván 2017; Tashjian and Galván 2020; Dimitrov et al. 2021), and provides preliminary evidence that habitually short sleepers may have a heightened sensitivity to time-on-task effects - mirroring prior results demonstrating that general time-on-task effects are exacerbated by experimental sleep loss (Asplund and Chee 2013; Hudson et al. 2020). Considering that the participants in our study largely had sleep durations within the recommended range for adults (7-8 hours), these results further imply that time-on-task analyses are sensitive to subtle changes in neural activity within a homogenous, healthy sample.

Self-reported and objective measures of sleep duration were differently associated with cognitive control activations and with task performance, providing additional evidence that self-reported versus objective measures of sleep - even within the same dimension - capture different phenomena (Klumpp et al. 2017; Bernstein et al. 2019; Stefansdottir et al. 2020; Scarlett et al. 2021; Tahmasian et al. 2021). Whereas the *objective* sleep duration was associated with time-on-task decreases in the anterior cingulate cortex - a core region for cognitive control - *self-reported* sleep duration was associated with time-on-task decreases in temporal areas, which are typically implicated in memory function or semantic processing. Also, objective sleep duration was most strongly associated with task performance (shorter duration associated with more variable RTs and more errors overall, as well as slower and more variable RTs with time on task). This supports extant findings linking shorter sleep to a more inattentive response style (Kuula et al. 2018; Saksvik-Lehouillier et al. 2020). Taken together, our results indicate that shorter objectively measured sleep duration is associated with poorer cognitive control functioning in normal sleepers, whereas the association between self-reported sleep duration and cognitive control function was less clear.

Through assessing the multidimensionality of habitual sleep health, some interesting overall patterns emerged. Most associations with cognitive control processing were related to later and more variable sleep timings. This provides additional evidence that sleep timing and variability are important aspects of sleep health (Chaput et al. 2020), and extends and substantiates prior studies indicating their relevance for brain functioning (Khalsa et al. 2016; Byrne et al. 2019; Facer-Childs et al. 2019; Lunsford-Avery et al. 2020; Zhang et al. 2020). Further, actigraphy-derived habitual sleep duration was the measure most closely associated with task performance, indicating that objective sleep duration is linked to performance-based cognitive control function in healthy young adults. For sleep efficiency and quality, cognitive control activations were significantly associated with levels of insomnia symptoms and actigraphy-derived sleep efficiency, but *not* with self-reported sleep quality (as measured using the PSQI). Whereas the ISI is specific to problems with going to sleep and maintaining sleep, the PSQI is a broader measure of perceived sleep quality (Chen et al. 2017). Our findings therefore imply that the efficiency and consistency of sleep, rather than perceived sleep quality, are more closely related to cognitive control function in normal sleepers.

The lack of a significant association with the PSQI is in contrast to several previous studies which have identified a link between this instrument and brain function (Elvemo et al. 2015; Curtis et al. 2016; Avinun et al. 2017; Klumpp et al. 2017; Cheng et al. 2018). However, most of these studies included patients or individuals with depressive symptoms, and / or were focused on emotion processing, and the PSQI has previously been shown to correlate with depression (Grandner et al. 2006; Klumpp et al. 2017). It is possible that poorer self-reported sleep quality, as measured using the PSQI, is more related to affective regulation (‘hot cognitive control function’) as compared to ‘cold’ cognitive processes (as was measured in our study) (Klumpp et al. 2017; Salehinejad et al. 2021). We also note that the participants in our study were considered eligible partly based on the *absence* of self-reported sleep and mental health problems, and hence, the overall prevalence of sleep problems was relatively low in our sample. These null-findings should thus be interpreted with caution. Future studies should continue to investigate how objective versus self-report measures of sleep relate to objective versus self-report measures of cognitive function, as well as mental health (Bernstein et al. 2019; Nicolazzo et al. 2021).

Due to the cross-sectional design of our study, we cannot conclude on the directionality of our results. Our findings may suggest that poorer sleep health leads to altered cognitive control function, or they may reflect inherent differences in cognitive control and/or brain function which give rise to real-life differences in sleep behavior. For example, those with a more reactive cognitive control style (i.e., a more ‘hyper-reactive’ brain signature) may be more impulsive in nature, leading to later and more variable bed/rise times. In line with this, previous studies have found a link between later chronotype preference and lower ability for self-regulation (Digdon and Howell 2008; Owens et al. 2016; Kuula et al. 2018). Also, circadian phenotype is known to be partially dependent on genetic factors (Katzenberg et al. 1998), which could in turn affect the development of cognitive control function and sleep behavior. On the other hand, it seems unlikely that cognitive control function would affect actigraphy-derived sleep efficiency, which was associated with increased neural recruitment in frontal areas with time on task. This finding in particular might reflect an increased need for cognitive compensation, or increased efforts to maintain alertness, as a result of insufficient sleep.

The restriction of inclusion to healthy, ‘normal sleepers’ between the ages of 20-40 ensured a homogenous age sample, and reduces the likelihood of results being confounded by other factors affecting sleep health (e.g., brain aging and health problems) (Scullin and Bliwise 2015; Bei et al. 2016; Alfini et al. 2020). This is an important strength of the current study. At the same time, it is important to acknowledge that the generalizability of our findings to other specific populations (e.g., other age cohorts, clinical populations, persons with significant sleep complaints, extreme chronotypes) - may be limited. Further, prior studies have shown that BOLD activations and task performance can vary throughout the day, and that people perform better when test times are aligned with their natural alertness peak (‘the synchrony effect’) (Goldstein et al. 2007; Schmidt et al. 2012; Song et al. 2018). One concern could therefore be that our results - in particular those pertaining to sleep timing and chronotype - were driven by inter-individual circadian effects. However, there was no systematic relationship between chronotype preference and test times in our study, and all testing was performed during standard working hours within a relatively large time-span (between 08:00 AM and 03:00 PM). There were also little-to-no associations between chronotype preference and task performance, except that later chronotype preference was associated with lower hit RT variability with increased time on task (Figure 4). As this study was not specifically designed to test circadian effects, conclusions regarding such effects are beyond the scope of the current study, and results should be interpreted with this in mind.

We collected a comprehensive selection of objective and self-reported measures in order to capture the multidimensionality of habitual sleep health. This can be considered both a strength and a limitation of our study. While the inclusion of different measures yields opportunity for replication and comparability of our results, it also introduces a considerable number of statistical tests with an increased risk of Type I errors. Our study has a reasonably large sample size (Poldrack et al. 2017), and is the largest study to date using neuroimaging combined with objective (actigraphy-based) assessment of habitual sleep health (Khalsa et al. 2016; Lunsford-Avery et al. 2020; Zhang et al. 2020). Statistical power may still be limited, as we are studying what is likely to be subtle effects. Additionally, each fMRI model was adjusted for age, sex, education, task performance, and head motion parameters in order to minimize the influence of potential confounding factors, which reduces the degrees of freedom. Some of our null-findings may therefore be Type-II errors as in similarly powered fMRI studies (Lieberman and Cunningham 2009).

To maintain a balance between the risk of Type 1 versus Type II errors, we did not formally adjust for multiple comparisons across the different fMRI GLMs (Rothman 1990; Lieberman and Cunningham 2009). The practical implementation of correcting p-values across different fMRI analyses based on our analytical approach is also not straight-forward (see Feise 2002; Lydersen 2021). However, each GLM was corrected for multiple comparisons using cluster-based inference, with a conventional cluster-defining threshold (Z = 3.1, corresponding to a p-value of .001) and a cluster probability threshold of p < .05 - determined using Gaussian RFT (Worsley 2011). This yielded relatively large and robust clusters (minimum cluster size across all analyses was 167 voxels) (Carp 2012; Woo et al. 2014). FSL’s FLAME has been shown to control the FWE rate relatively strictly at this cluster-defining threshold as compared to other tools (Eklund et al. 2016), and, generally, using more stringent cluster thresholds increases the risk of Type II errors substantially (Woo et al. 2014). The distinction between proactive and reactive cognitive control processing, as well as associated time-on-task effects, is well grounded in previous literature, and clearly operationalized and implemented in our well-validated fMRI protocol (Olsen et al. 2013). Our overarching interpretation of results, which is based on multiple statistically significant findings (i.e., patterns of findings), is therefore supported by a clear theoretical framework, and provides important groundwork for future testing and/or replication.

In conclusion, in this study of adult, normal sleepers, we found that poorer sleep health was associated with a hyper-reactive brain state during a test of cognitive control function, as well as increased proactive cognitive control processing with longer time on task. Across the different dimensions of sleep health, later and more variable sleep timing was most closely associated with higher cognitive control BOLD activations, whereas shorter objective sleep duration was associated with poorer task performance and lower BOLD activations with time on task. Given that our fMRI analyses were adjusted for performance, we suggest that the altered brain activity observed with poorer sleep health may reflect compensatory neural recruitment and / or inefficient neural processing. Future studies should continue to focus on naturalistic measurement of normal sleep to elucidate the complex relationships between sleep and brain function.

## Competing Interest Statements

PMT received partial grant support from Biogen, Inc., for research unrelated to this manuscript. AO is an owner and medical advisor for Nordic Brain Tech AS.

## Funding

This work was supported by the Liaison Committee between the Central Norway Regional Health Authority (RHA) and the Norwegian University of Science and Technology (NTNU) (2020/39645) to AO.

## Acknowledgments

We thank all the participants in the study. We thank Ingeborg Nakken for her assistance in optimizing and implementing the MRI protocol. We also thank Anne Margrethe Berggrav, Ella Flagstad, Tea Høiby, Sofie Jakobsen, Ole Erik Melsæter, Marte Sandbukt Petterson, Marte Hofstad, Adrian Helgå Vestøl for their contributions to participant recruitment and data collection. PMT is supported in part by the National Institutes of Health (R01MH116147).

## Notes

### Competing Interest Statement

The authors have declared no competing interest.

### Summary of Updates

Revised according to reviewer comments.

## References

Abraham A. 2013. The World According to Me: Personal Relevance and the Medial Prefrontal Cortex. Front Hum Neurosci. 7.

Akerstedt T, Gillberg M. 1990. Subjective and objective sleepiness in the active individual. Int J Neurosci. 52:29–37.

Alfini AJ, Tzuang M, Owusu JT, Spira AP. 2020. Later-life sleep, cognition, and neuroimaging research: an update for 2020. Curr Opin Behav Sci, Sleep and cognition. 33:72–77.

Allen S, Elder G, Longstaff L, Gotts Z, Sharman R, Akram U, Ellis J. 2018. Exploration of Potential Objective and Subjective Indicators of Sleep Health in Normal Sleepers. Nat Sci Sleep. 10:303–312.

Almutairi B, Langley C, Crawley E, Thai NJ. 2020. Using structural and functional MRI as a neuroimaging technique to investigate chronic fatigue syndrome/myalgic encephalopathy: a systematic review. BMJ Open. 10:e031672.

Andersson JLR, Jenkinson M, Smith S. 2007. Non-linear registration aka Spatial normalisation (Technical Report No. TR07JA2). FMRIB Centre, Oxford, United Kingdom.

Andersson JLR, Skare S, Ashburner J. 2003. How to correct susceptibility distortions in spin-echo echo-planar images: application to diffusion tensor imaging. NeuroImage. 20:870–888.

Asplund CL, Chee MWL. 2013. Time-on-task and sleep deprivation effects are evidenced in overlapping brain areas. NeuroImage. 82:326–335.

Avants BB, Epstein CL, Grossman M, Gee JC. 2008. Symmetric diffeomorphic image registration with cross-correlation: Evaluating automated labeling of elderly and neurodegenerative brain. Med Image Anal, Special Issue on The Third International Workshop on Biomedical Image Registration – WBIR 2006. 12:26–41.

Avinun R, Nevo A, Knodt AR, Elliott ML, Radtke SR, Brigidi BD, Hariri AR. 2017. Reward-Related Ventral Striatum Activity Buffers against the Experience of Depressive Symptoms Associated with Sleep Disturbances. J Neurosci. 37:9724–9729.

Badre D. 2008. Cognitive control, hierarchy, and the rostro-caudal organization of the frontal lobes. Trends Cogn Sci. 12:193–200.

Beattie L, Espie CA, Kyle SD, Biello SM. 2015. How are normal sleeping controls selected? A systematic review of cross-sectional insomnia studies and a standardized method to select healthy controls for sleep research. Sleep Med. 16:669–677.

Bei B, Wiley JF, Trinder J, Manber R. 2016. Beyond the mean: A systematic review on the correlates of daily intraindividual variability of sleep/wake patterns. Sleep Med Rev. 28:108–124.

Bernstein JPK, DeVito A, Calamia M. 2019. Subjectively and Objectively Measured Sleep Predict Differing Aspects of Cognitive Functioning in Adults. Arch Clin Neuropsychol. 34:1127–1137.

Bliwise DL, Young TB. 2007. The Parable of Parabola: What the U-Shaped Curve Can and Cannot Tell Us about Sleep. Sleep. 30:1614–1615.

Boyne K, Sherry DD, Gallagher PR, Olsen M, Brooks LJ. 2013. Accuracy of computer algorithms and the human eye in scoring actigraphy. Sleep Breath. 17:411–417.

Braver TS. 2012. The variable nature of cognitive control: A dual-mechanisms framework. Trends Cogn Sci. 16:106–113.

Burgess GC, Braver TS. 2010. Neural mechanisms of interference control in working memory: Effects of interference expectancy and fluid intelligence. PLoS ONE. 5:1–11.

Buysse DJ. 2014. Sleep health: can we define it? Does it matter? Sleep. 37:9–17.

Buysse DJ, Hall ML, Strollo PJ, Kamarck TW, Owens J, Lee L, Reis SE, Matthews KA. 2008. Relationships Between the Pittsburgh Sleep Quality Index (PSQI), Epworth Sleepiness Scale (ESS), and Clinical/Polysomnographic Measures in a Community Sample. J Clin Sleep Med JCSM Off Publ Am Acad Sleep Med. 4:563.

Buysse DJ, Reynolds CF, Monk TH, Hoch CC, Yeager A, Kupfer DJ. 1991. Quantification of subjective sleep quality in healthy elderly men and women using the Pittsburgh Sleep Quality Index (PSQI). Sleep. 14:331–338.

Byrne JEM, Tremain H, Leitan ND, Keating C, Johnson SL, Murray G. 2019. Circadian modulation of human reward function: Is there an evidentiary signal in existing neuroimaging studies? Neurosci Biobehav Rev. 99:251–274.

Cai W, Chen T, Ryali S, Kochalka J, Li C-SR, Menon V. 2016. Causal Interactions Within a Frontal-Cingulate-Parietal Network During Cognitive Control: Convergent Evidence from a Multisite–Multitask Investigation. Cereb Cortex. 26:2140–2153.

Carney CE, Buysse DJ, Ancoli-Israel S, Edinger JD, Krystal AD, Lichstein KL, Morin CM. 2012. The Consensus Sleep Diary: Standardizing Prospective Sleep Self-Monitoring. Sleep. 35:287–302.

Carp J. 2012. The secret lives of experiments: Methods reporting in the fMRI literature. NeuroImage. 63:289–300.

Chalder T, Berelowitz G, Pawlikowska T, Watts L, Wessely S, Wright D, Wallace EP. 1993. Development of a fatigue scale. J Psychosom Res.

Chaput J-P, Dutil C, Featherstone R, Ross R, Giangregorio L, Saunders TJ, Janssen I, Poitras VJ, Kho ME, Ross-White A, Zankar S, Carrier J. 2020. Sleep timing, sleep consistency, and health in adults: a systematic review. Appl Physiol Nutr Metab. 45:S232–S247.

Chee MWL, Tan JC. 2010. Lapsing when sleep deprived: Neural activation characteristics of resistant and vulnerable individuals. NeuroImage. 51:835–843.

Chen P-Y, Jan Y-W, Yang C-M. 2017. Are the Insomnia Severity Index and Pittsburgh Sleep Quality Index valid outcome measures for Cognitive Behavioral Therapy for Insomnia? Inquiry from the perspective of response shifts and longitudinal measurement invariance in their Chinese versions. Sleep Med. 35:35–40.

Cheng W, Rolls E, Gong W, Du J, Zhang J, Zhang X-Y, Li F, Feng J. 2020. Sleep duration, brain structure, and psychiatric and cognitive problems in children. Mol Psychiatry.

Cheng W, Rolls ET, Ruan H, Feng J. 2018. Functional Connectivities in the Brain That Mediate the Association between Depressive Problems and Sleep Quality. JAMA Psychiatry. 75:1052–1061.

Chevalier N, Martis SB, Curran T, Munakata Y. 2015. Metacognitive Processes in Executive Control Development: The Case of Reactive and Proactive Control. J Cogn Neurosci. 27:1125–1136.

Cole MW, Repovš G, Anticevic A. 2014. The frontoparietal control system: a central role in mental health. Neurosci Rev J Bringing Neurobiol Neurol Psychiatry. 20:652–664.

Conners CK. 2014. Conners’ Continuous Performance Test 3rd edition manual. 3rd ed. Toronto, CA: Multi-Health Systems.

Cook DB, O’Connor PJ, Lange G, Steffener J, O’Connor PJ, Lange G, Steffener J. 2007. Functional neuroimaging correlates of mental fatigue induced by cognition among chronic fatigue syndrome patients and controls. NeuroImage. 36:108–122.

Curtis BJ, Williams PG, Jones CR, Anderson JS. 2016. Sleep duration and resting fMRI functional connectivity: examination of short sleepers with and without perceived daytime dysfunction. Brain Behav. 6:1–13.

Dale AM, Fischl B, Sereno MI. 1999. Cortical Surface-Based Analysis: I. Segmentation and Surface Reconstruction. NeuroImage. 9:179–194.

DeLuca J, Genova HM, Capili EJ, Wylie GR. 2009. Functional Neuroimaging of Fatigue. Phys Med Rehabil Clin N Am. 20:325–337.

Diamond A. 2013. Executive functions. Annu Rev Psychol. 64:135–168.

Digdon NL, Howell AJ. 2008. College students who have an eveningness preference report lower self-control and greater procrastination. Chronobiol Int. 25:1029–1046.

Dimitrov A, Nowak J, Ligdorf A, Oei NYL, Adli M, Walter H, Veer IM. 2021. Natural sleep loss is associated with lower mPFC activity during negative distracter processing. Cogn Affect Behav Neurosci. 21:242–253.

Dong L, Martinez AJ, Buysse DJ, Harvey AG. 2019. A composite measure of sleep health predicts concurrent mental and physical health outcomes in adolescents prone to eveningness. Sleep Health. 5:166–174.

Dosenbach NUF, Fair DA, Cohen AL, Schlaggar BL, Petersen SE. 2008. A dual-networks architecture of top-down control. Trends Cogn Sci. 12:99–105.

Dosenbach NUF, Visscher KM, Palmer ED, Miezin FM, Wenger KK, Kang HC, Burgund ED, Grimes AL, Schlaggar BL, Petersen SE. 2006. A Core System for the Implementation of Task Sets. Neuron. 50:799–812.

Drummond SPA, Meloy MJ, Yanagi MA, Orff HJ, Brown GG. 2005. Compensatory recruitment after sleep deprivation and the relationship with performance. Psychiatry Res - Neuroimaging. 140:211–223.

Edinger JD, Bonnet MH, Bootzin RR, Doghramji K, Dorsey CM, Espie CA, Jamieson AO, McCall WV, Morin CM, Stepanski EJ. 2004. Derivation of research diagnostic criteria for insomnia: Report of an American Academy of Sleep Medicine work group. Sleep. 27:1567–1596.

Eklund A, Nichols TE, Knutsson H. 2016. Cluster failure: Why fMRI inferences for spatial extent have inflated false-positive rates. Proc Natl Acad Sci U S A. 113:7900–7905.

Elvemo NA, Landrø NI, Borchgrevink PC, Håberg AK. 2015. A particular effect of sleep, but not pain or depression, on the blood-oxygen-level dependent response during working memory tasks in patients with chronic pain. J Pain Res. 8:335–346.

Esteban O, Blair RW, Markiewicz CJ, Berleant SL, Moodie CA, Ma F, Isik AI, Erramuzpe A, Kent JD, Goncalves M, DuPre E, Sitek KR, Gomez DEP, Lurie DJ, Ye Z, Salo T, Valabregue R, Amlien IK, Liem F, Jacoby N, Stojic H, Halchenko YO, Rivera-Dompenciel A, Ciric R, Sneve MH, Heinsfeld AS, Thompson WH, Tooley UA, Poldrack RA, Gorgolewski KJ. 2018. fMRIPrep.

Esteban O, Markiewicz CJ, Blair RW, Moodie CA, Isik AI, Erramuzpe A, Kent JD, Goncalves M, DuPre E, Snyder M, Oya H, Ghosh SS, Wright J, Durnez J, Poldrack RA, Gorgolewski KJ. 2019. fMRIPrep: a robust preprocessing pipeline for functional MRI. Nat Methods. 16:111–116.

Etkin A, Egner T, Kalisch R. 2011. Emotional processing in anterior cingulate and medial prefrontal cortex. Trends Cogn Sci. 15:85–93.

Evans MA, Buysse DJ, Marsland AL, Wright AGC, Foust J, Carroll LW, Kohli N, Mehra R, Jasper A, Srinivasan S, Hall MH. 2021. Meta-analysis of age and actigraphy-assessed sleep characteristics across the lifespan. Sleep.

Faaland P, Vedaa Ø, Langsrud K, Sivertsen B, Lydersen S, Vestergaard CL, Kjørstad K, Vethe D, Ritterband LM, Harvey AG, Stiles TC, Scott J, Kallestad H. 2022. Digital cognitive behaviour therapy for insomnia (dCBT-I): Chronotype moderation on intervention outcomes. J Sleep Res. 31:e13572.

Facer-Childs ER, Campos BM, Middleton B, Skene DJ, Bagshaw AP. 2019. Circadian phenotype impacts the brain’s resting-state functional connectivity, attentional performance, and sleepiness. Sleep. 42.

Fales CL, Barch DM, Burgess GC, Schaeffer A, Mennin DS, Gray JR, Braver TS. 2008. Anxiety and cognitive efficiency: Differential modulation of transient and sustained neural activity during a working memory task. Cogn Affect Behav Neurosci. 8:239–253.

Fasiello E, Gorgoni M, Scarpelli S, Alfonsi V, Ferini Strambi L, De Gennaro L. 2022. Functional connectivity changes in insomnia disorder: A systematic review. Sleep Med Rev. 61:101569.

Feise RJ. 2002. Do multiple outcome measures require p-value adjustment? BMC Med Res Methodol. 2:8.

Follesø HS, Austad SB, Olsen A, Saksvik-Lehouillier I. 2021. The development, inter-rater agreement and performance of a hierarchical procedure for setting the rest-interval in actigraphy data. Sleep Med. 85:221–229.

Freeman D, Sheaves B, Waite F, Harvey AG, Harrison PJ. 2020. Sleep disturbance and psychiatric disorders. Lancet Psychiatry. 7:628–637.

Fueggle SN, Bucks RS, Fox AM. 2018. The relationship between naturalistic sleep variation and error monitoring in young adults: An event-related potential (ERP) study. Int J Psychophysiol. 134:151–158.

Goldstein D, Hahn CS, Hasher L, Wiprzycka UJ, Zelazo PD. 2007. Time of day, intellectual performance, and behavioral problems in Morning versus Evening type adolescents: Is there a synchrony effect? Personal Individ Differ. 42:431–440.

Gorgolewski K, Burns C, Madison C, Clark D, Halchenko Y, Waskom M, Ghosh S. 2011. Nipype: A Flexible, Lightweight and Extensible Neuroimaging Data Processing Framework in Python. Front Neuroinformatics. 5.

Gorgolewski KJ, Esteban O, Markiewicz CJ, Ziegler E, Ellis DG, Notter MP, Jarecka D, et al. 2018. Nipype.

Goschke T. 2013. Dysfunctions of decision-making and cognitive control as transdiagnostic mechanisms of mental disorders: Advances, gaps, and needs in current research. Int J Methods Psychiatr Res. 23:41–57.

Grandner MA, Kripke DF, Yoon IY, Youngstedt SD. 2006. Criterion validity of the Pittsburgh Sleep Quality Index: Investigation in a non-clinical sample. Sleep Biol Rhythms. 4:129–136.

Grinband J, Savitskaya J, Wager TD, Teichert T, Ferrera VP, Hirsch J. 2011. The dorsal medial frontal cortex is sensitive to time on task, not response conflict or error likelihood. NeuroImage. 57:303–311.

Grolemund G, Wickham H. 2011. Dates and Times Made Easy with lubridate. J Stat Softw. 40:1–25.

Grumbach P, Opel N, Martin S, Meinert S, Leehr EJ, Redlich R, Enneking V, Goltermann J, Baune BT, Dannlowski U, Repple J. 2020. Sleep duration is associated with white matter microstructure and cognitive performance in healthy adults. Hum Brain Mapp. 41:4397–4405.

Guadagni V, Burles F, Ferrara M, Iaria G. 2018. Sleep quality and its association with the insular cortex in emotional empathy. Eur J Neurosci. 48:2288–2300.

Gurvich C, Hoy K, Thomas N, Kulkarni J. 2018. Sex Differences and the Influence of Sex Hormones on Cognition through Adulthood and the Aging Process. Brain Sci. 8:163.

Hershner S. 2020. Sleep and academic performance: measuring the impact of sleep. Curr Opin Behav Sci, Sleep and cognition. 33:51–56.

Hirshkowitz M, Whiton K, Albert SM, Alessi C, Bruni O, DonCarlos L, Hazen N, Herman J, Katz ES, Kheirandish-Gozal L, Neubauer DN, O’Donnell AE, Ohayon M, Peever J, Rawding R, Sachdeva RC, Setters B, Vitiello MV, Ware JC, Adams Hillard PJ. 2015. National sleep foundation’s sleep time duration recommendations: Methodology and results summary. Sleep Health. 1:40–43.

Hokett E, Arunmozhi A, Campbell J, Verhaeghen P, Duarte A. 2021. A systematic review and meta-analysis of individual differences in naturalistic sleep quality and episodic memory performance in young and older adults. Neurosci Biobehav Rev. 127:675–688.

Horne JA, Ostberg O. 1976. A self-assessment questionnaire to determine morningness-eveningness in human circadian rhythms. Int J Chronobiol. 4:97–110.

Huang S, Zhu Z, Zhang W, Chen Y, Zhen S. 2017. Trait impulsivity components correlate differently with proactive and reactive control. PLOS ONE. 12:e0176102.

Hudson AN, Van Dongen HPA, Honn KA. 2020. Sleep deprivation, vigilant attention, and brain function: a review. Neuropsychopharmacology. 45:21–30.

Husa RA, Buchanan TW, Kirchhoff BA. 2022. Subjective stress and proactive and reactive cognitive control strategies. Eur J Neurosci. 55:2558–2570.

Itani O, Jike M, Watanabe N, Kaneita Y. 2017. Short sleep duration and health outcomes: a systematic review, meta-analysis, and meta-regression. Sleep Med. 32:246–256.

Jenkins AC, Mitchell JP. 2011. Medial prefrontal cortex subserves diverse forms of self-reflection. Soc Neurosci. 6:211–218.

Jenkinson M, Bannister P, Brady M, Smith S. 2002. Improved optimization for the robust and accurate linear registration and motion correction of brain images. NeuroImage. 17:825–841.

Jenkinson M, Beckmann CF, Behrens TEJ, Woolrich MW, Smith SM. 2012. FSL. NeuroImage. 62:782–790.

Jenkinson M, Smith S. 2001. A global optimisation method for robust affine registration of brain images. Med Image Anal. 5:143–156.

Johns MW. 1991. A New Method for Measuring Daytime Sleepiness: The Epworth Sleepiness Scale. Am Sleep Disord Assoc Sleep Res Soc. 14:540–545.

Kato K, Iwamoto K, Kawano N, Noda Y, Ozaki N, Noda A. 2018. Differential effects of physical activity and sleep duration on cognitive function in young adults. J Sport Health Sci. 7:227–236.

Katzenberg D, Young T, Finn L, Lin L, King DP, Takahashi JS, Mignot E. 1998. A CLOCK Polymorphism Associated with Human Diurnal Preference. Sleep. 21:569–576.

Khalsa S, Hale JR, Goldstone A, Wilson RS, Mayhew SD, Bagary M, Bagshaw AP. 2017. Habitual sleep durations and subjective sleep quality predict white matter differences in the human brain. Neurobiol Sleep Circadian Rhythms. 3:17–25.

Khalsa S, Mayhew SD, Przezdzik I, Wilson R, Hale J, Goldstone A, Bagary M, Bagshaw AP. 2016. Variability in Cumulative Habitual Sleep Duration Predicts Waking Functional Connectivity. Sleep. 39:87–95.

Kim REY, Abbott RD, Kim S, Thomas RJ, Yun C-H, Kim H, Johnson H, Shin C. 2021. Sleep Duration, Sleep Apnea, and Gray Matter Volume. J Geriatr Psychiatry Neurol. 891988720988918.

Kleerekooper I, van Rooij SJH, van den Wildenberg WPM, de Leeuw M, Kahn RS, Vink M. 2016. The effect of aging on fronto-striatal reactive and proactive inhibitory control. NeuroImage. 132:51–58.

Klein A, Ghosh SS, Bao FS, Giard J, Häme Y, Stavsky E, Lee N, Rossa B, Reuter M, Neto EC, Keshavan A. 2017. Mindboggling morphometry of human brains. PLOS Comput Biol. 13:e1005350.

Kluger BM, Krupp LB, Enoka RM. 2013. Fatigue and fatigability in neurologic illnesses: Proposal for a unified taxonomy. Neurology. 80:409–416.

Klumpp H, Roberts J, Kapella MC, Kennedy AE, Kumar A, Phan KL. 2017. Subjective and objective sleep quality modulate emotion regulatory brain function in anxiety and depression. Depress Anxiety. 34:651–660.

Knutson KL, von Schantz M. 2018. Associations between chronotype, morbidity and mortality in the UK Biobank cohort. Chronobiol Int. 35:1045–1053.

Kohl AD, Wylie GR, Genova HM, Hillary FG, DeLuca J. 2009. The neural correlates of cognitive fatigue in traumatic brain injury using functional MRI. Brain Inj. 23:420–432.

Krause AJ, Simon EB, Mander BA, Greer SM, Saletin JM, Goldstein-Piekarski AN, Walker MP. 2017. The sleep-deprived human brain. Nat Rev Neurosci. 18:404–418.

Kubota M, Hadley LV, Schaeffner S, Könen T, Meaney J-A, Auyeung B, Morey CC, Karbach J, Chevalier N. 2020. Consistent use of proactive control and relation with academic achievement in childhood. Cognition. 203:104329.

Kuula L, Pesonen AK, Heinonen K, Kajantie E, Eriksson JG, Andersson S, Lano A, Lahti J, Wolke D, Räikkönen K. 2018. Naturally occurring circadian rhythm and sleep duration are related to executive functions in early adulthood. J Sleep Res. 27:113–119.

Landry GJ, Best JR, Liu-Ambrose T. 2015. Measuring sleep quality in older adults: a comparison using subjective and objective methods. Front Aging Neurosci. 7:166.

Langner R, Eickhoff SB. 2013. Sustaining attention to simple tasks: A meta-analytic review of the neural mechanisms of vigilant attention. Psychol Bull. 139:870–900.

Lauderdale DS, Knutson KL, Yan LL, Liu K, Rathouz PJ. 2008. Sleep duration: how well do self-reports reflect objective measures? The CARDIA Sleep Study. Epidemiol Camb Mass. 19:838–845.

Lavie P. 2009. Self-reported sleep duration – what does it mean? J Sleep Res. 18:385–386.

Lesh TA, Westphal AJ, Niendam TA, Yoon JH, Minzenberg MJ, Ragland JD, Solomon M, Carter CS. 2013. Proactive and reactive cognitive control and dorsolateral prefrontal cortex dysfunction in first episode schizophrenia. NeuroImage Clin. 2:590–599.

Lieberman MD, Cunningham WA. 2009. Type I and Type II error concerns in fMRI research: re-balancing the scale. Soc Cogn Affect Neurosci. 4:423–428.

Lim J, Dinges DF. 2010. A meta-analysis of the impact of short-term sleep deprivation on cognitive variables. Psychol Bull. 136:375–389.

Lim J, Wu W-C, Wang J, Detre JA, Dinges DF, Rao H. 2010. Imaging brain fatigue from sustained mental workload: An ASL perfusion study of the time-on-task effect. NeuroImage. 49:3426–3435.

Lowe CJ, Safati A, Hall PA. 2017. The neurocognitive consequences of sleep restriction: A meta-analytic review. Neurosci Biobehav Rev. 80:586–604.

Lunsford-Avery JR, Damme KSF, Engelhard MM, Kollins SH, Mittal VA. 2020. Sleep/Wake Regularity Associated with Default Mode Network Structure among Healthy Adolescents and Young Adults. Sci Rep. 10:1–8.

Lydersen S. 2021. Adjustment of p-values for multiple hypotheses. Tidsskr Den Nor Laegeforening Tidsskr Prakt Med Ny Raekke. 141.

Maire M, Reichert CF, Gabel V, Viola AU, Phillips C, Berthomier C, Borgwardt S, Cajochen C, Schmidt C. 2018. Human brain patterns underlying vigilant attention: Impact of sleep debt, circadian phase and attentional engagement. Sci Rep. 8:970.

Makowski D. 2018. The psycho Package: An Efficient and Publishing-Oriented Workflow for Psychological Science. J Open Source Softw. 3:470.

Mantua J, Simonelli G. 2019. Sleep duration and cognition: is there an ideal amount? Sleep. 42:1–3.

Matthews KA, Patel SR, Pantesco EJ, Buysse DJ, Kamarck TW, Lee L, Hall MH. 2018. Similarities and differences in estimates of sleep duration by polysomnography, actigraphy, diary, and self-reported habitual sleep in a community sample. Sleep Health. 4:96–103.

McSorley VE, Bin YS, Lauderdale DS. 2019. Associations of Sleep Characteristics With Cognitive Function and Decline Among Older Adults. Am J Epidemiol. 188:1066–1075.

McTeague LM, Goodkind MS, Etkin A. 2016. Transdiagnostic impairment of cognitive control in mental illness. J Psychiatr Res. 83:37–46.

Menon V, Gallardo G, Pinsk MA, Nguyen V-D, Li J-R, Cai W, Wassermann D. 2020. Microstructural organization of human insula is linked to its macrofunctional circuitry and predicts cognitive control. eLife. 9:e53470.

Mignot E. 2008. Why We Sleep: The Temporal Organization of Recovery. PLOS Biol. 6:e106.

Morin CM. 1993. Insomnia: Psychological Assessment and Management. New York, NY: Guildford Press.

Neubauer AC, Fink A. 2009. Intelligence and neural efficiency. Neurosci Biobehav Rev. 33:1004–1023.

Nicolazzo J, Xu K, Lavale A, Buckley R, Yassi N, Hamilton GS, Maruff P, Baril A-A, Lim YY, Pase MP. 2021. Sleep symptomatology is associated with greater subjective cognitive concerns: Findings from the community-based Healthy Brain Project. Sleep.

Niebaum JC, Chevalier N, Guild RM, Munakata Y. 2020. Developing adaptive control: Age-related differences in task choices and awareness of proactive and reactive control demands. Cogn Affect Behav Neurosci.

Niendam TA, Laird AR, Ray KL, Dean YM, Glahn DC, Carter CS. 2012. Meta-analytic evidence for a superordinate cognitive control network subserving diverse executive functions. Cogn Affect Behav Neurosci. 12:241–268.

O’Doherty JP. 2004. Reward representations and reward-related learning in the human brain: insights from neuroimaging. Curr Opin Neurobiol. 14:769–776.

Ohayon M, Wickwire EM, Hirshkowitz M, Albert SM, Avidan A, Daly FJ, Dauvilliers Y, Ferri R, Fung C, Gozal D, Hazen N, Krystal A, Lichstein K, Mallampalli M, Plazzi G, Rawding R, Scheer FA, Somers V, Vitiello MV. 2017. National Sleep Foundation’s sleep quality recommendations: first report. Sleep Health. 3:6–19.

Oldfield RC. 1971. The assessment and analysis of handedness: The Edinburgh inventory. Neuropsychologia.

Olsen A, Babikian T, Dennis EL, Ellis-Blied MU, Giza C, Marion SD, Mink R, Johnson J, Babbitt CJ, Thompson PM, Asarnow RF. 2020. Functional Brain Hyperactivations Are Linked to an Electrophysiological Measure of Slow Interhemispheric Transfer Time after Pediatric Moderate/Severe Traumatic Brain Injury. J Neurotrauma. 37:397–409.

Olsen A, Brunner JF, Indredavik Evensen KA, Finnanger TG, Vik A, Skandsen T, Landrø NI, Håberg AK. 2015. Altered Cognitive Control Activations after Moderate-to-Severe Traumatic Brain Injury and Their Relationship to Injury Severity and Everyday-Life Function. Cereb Cortex. 25:2170–2180.

Olsen A, Dennis EL, Evensen KAI, Husby Hollund IM, Løhaugen GCC, Thompson PM, Brubakk A-M, Eikenes L, Håberg AK. 2018. Preterm birth leads to hyper-reactive cognitive control processing and poor white matter organization in adulthood. NeuroImage. 167:419–428.

Olsen A, Ferenc Brunner J, Evensen KAI, Garzon B, Landrø NI, Håberg AK. 2013. The Functional Topography and Temporal Dynamics of Overlapping and Distinct Brain Activations for Adaptive Task Control and Stable Task-set Maintenance during Performance of an fMRI-adapted Clinical Continuous Performance Test. J Cogn Neurosci. 25:903–919.

Owens JA, Dearth-Wesley T, Lewin D, Gioia G, Whitaker RC. 2016. Self-Regulation and Sleep Duration, Sleepiness, and Chronotype in Adolescents. Pediatrics. 138.

Palmer CA, Alfano CA. 2017. Sleep and emotion regulation: An organizing, integrative review. Sleep Med Rev. 31:6–16.

Petersen SE, Dubis JW. 2012. The mixed block/event-related design. NeuroImage. 62:1177–1184.

Petersen SE, van Mier H, Fiez JA, Raichle MA. 1998. The effects of practice on the functional anatomy of task performance. Proc Natl Acad Sci. 98:853–860.

Poldrack RA, Baker CI, Durnez J, Gorgolewski KJ, Matthews PM, Munafò MR, Nichols TE, Poline J-B, Vul E, Yarkoni T. 2017. Scanning the horizon: towards transparent and reproducible neuroimaging research. Nat Rev Neurosci. 18:115–126.

Price CJ, Crinion J, Friston KJ. 2006. Design and analysis of fMRI studies with neurologically impaired patients. J Magn Reson Imaging. 23:816–826.

Qin S, Leong RLF, Ong JL, Chee MWL. 2023. Associations between objectively measured sleep parameters and cognition in healthy older adults: A meta-analysis. Sleep Med Rev. 67:101734.

Revelle W. 2022. psych: Procedures for Psychological, Psychometric, and Personality Research.

Richards A, Inslicht SS, Metzler TJ, Mohlenhoff BS, Rao MN, O’Donovan A, Neylan TC. 2017. Sleep and Cognitive Performance From Teens To Old Age: More Is Not Better. Sleep. 40.

Riemann D, Spiegelhalder K, Feige B, Voderholzer U, Berger M, Perlis M, Nissen C. 2010. The hyperarousal model of insomnia: A review of the concept and its evidence. Sleep Med Rev. 14:19–31.

Robbins R, Quan SF, Barger LK, Czeisler CA, Fray-Witzer M, Weaver MD, Zhang Y, Redline S, Klerman EB. 2021. Self-reported sleep duration and timing: A methodological review of event definitions, context, and timeframe of related questions. Sleep Epidemiol. 1:100016.

Rothman KJ. 1990. No Adjustments Are Needed for Multiple Comparisons. Epidemiology. 1:43–46.

Saksvik-Lehouillier I, Saksvik SB, Dahlberg J, Tanum TK, Ringen H, Karlsen HR, Smedbøl T, Sørengaard TA, Stople M, Kallestad H, Olsen A. 2020. Mild to moderate partial sleep deprivation is associated with increased impulsivity and decreased positive affect in young adults. Sleep. 43.

Salehinejad MA, Ghanavati E, Rashid MHA, Nitsche MA. 2021. Hot and cold executive functions in the brain: A prefrontal-cingular network. Brain Neurosci Adv. 5:23982128211007770.

Satterfield BC, Wisor JP, Schmidt MA, Van Dongen HPA. 2017. Time-on-Task Effect During Sleep Deprivation in Healthy Young Adults Is Modulated by Dopamine Transporter Genotype. SLEEP. 40.

Scarlett S, Nolan HN, Kenny RA, O’Connell MDL. 2021. Discrepancies in self-reported and actigraphy-based sleep duration are associated with self-reported insomnia symptoms in community-dwelling older adults. Sleep Health. 7:83–92.

Schiel JE, Tamm S, Holub F, Petri R, Dashti HS, Domschke K, Feige B, Lane JM, Riemann D, Rutter MK, Saxena R, Tahmasian M, Wang H, Kyle SD, Spiegelhalder K. 2022. Associations Between Sleep Health and Amygdala Reactivity to Negative Facial Expressions in the UK Biobank Cohort. Biol Psychiatry, Threat, Stress, and Health. 92:693–700.

Schmid PC, Kleiman T, Amodio DM. 2015. Neural mechanisms of proactive and reactive cognitive control in social anxiety. Cortex. 70:137–145.

Schmidt C, Collette F, Reichert CF, Maire M, Vandewalle G, Peigneux P, Cajochen C. 2015. Pushing the Limits: Chronotype and Time of Day Modulate Working Memory-Dependent Cerebral Activity. Front Neurol. 6:199.

Schmidt C, Peigneux P, Leclercq Y, Sterpenich V, Vandewalle G, Phillips C, Berthomier P, Berthomier C, Tinguely G, Gais S, Schabus M, Desseilles M, Dang-Vu T, Salmon E, Degueldre C, Balteau E, Luxen A, Cajochen C, Maquet P, Collette F. 2012. Circadian preference modulates the neural substrate of conflict processing across the day. PloS One. 7:e29658.

Scullin MK, Bliwise DL. 2015. Sleep, Cognition, and Normal Aging: Integrating a Half Century of Multidisciplinary Research. Perspect Psychol Sci. 10:97–137.

Seeley WW, Menon V, Schatzberg AF, Keller J, Glover GH, Kenna H, Reiss AL, Greicius MD. 2007. Dissociable Intrinsic Connectivity Networks for Salience Processing and Executive Control. J Neurosci. 27:2349–2356.

Sexton CE, Storsve AB, Walhovd KB, Johansen-Berg H, Fjell AM. 2014. Poor sleep quality is associated with increased cortical atrophy in community-dwelling adults. Neurology. 83:967–973.

Shahid A, Wilkinson K, Marcu S, Shapiro CM. 2011. Karolinska Sleepiness Scale (KSS). In: Shahid A,, Wilkinson K,, Marcu S,, Shapiro CM, editors. STOP, THAT and One Hundred Other Sleep Scales. New York, NY: Springer New York. p. 209–210.

Shanmugan S, Satterthwaite TD. 2016. Neural markers of the development of executive function: relevance for education. Curr Opin Behav Sci, Neuroscience of education. 10:7–13.

Smith SM. 2002. Fast robust automated brain extraction. Hum Brain Mapp. 17:143–155.

Snyder HR, Miyake A, Hankin BL. 2015. Advancing understanding of executive function impairments and psychopathology: Bridging the gap between clinical and cognitive approaches. Front Psychol. 6.

Song J, Feng P, Zhao X, Xu W, Xiao L, Zhou J, Zheng Y. 2018. Chronotype regulates the neural basis of response inhibition during the daytime. Chronobiol Int. 35:208–218.

Staub B, Doignon-Camus N, Bacon É, Bonnefond A. 2014. Age-related differences in the recruitment of proactive and reactive control in a situation of sustained attention. Biol Psychol. 103:38–47.

Stefansdottir R, Gundersen H, Rognvaldsdottir V, Lundervold AS, Gestsdottir S, Gudmundsdottir SL, Chen KY, Brychta RJ, Johannsson E. 2020. Association between free-living sleep and memory and attention in healthy adolescents. Sci Rep. 10:16877.

Stefansdottir R, Rognvaldsdottir V, Chen KY, Johannsson E, Brychta RJ. 2022. Sleep timing and consistency are associated with the standardised test performance of Icelandic adolescents. J Sleep Res. 31:e13422.

Suemoto CK, Santos RB, Giatti S, Aielo AN, Silva WA, Parise BK, Cunha LF, Souza SP, Griep RH, Brunoni AR, Lotufo PA, Bensenor IM, Drager LF. 2022. Association between objective sleep measures and cognitive performance: a cross-sectional analysis in the Brazilian Longitudinal Study of Adult Health (ELSA-Brasil) study. J Sleep Res. n/a:e13659.

Tahmasian M, Aleman A, Andreassen OA, Arab Z, Baillet M, Benedetti F, Bresser T, Bright J, Chee MWL, Chylinski D, Cheng W, Deantoni M, Dresler M, Eickhoff SB, Eickhoff CR, Elvsåshagen T, Feng J, Foster-Dingley JC, Ganjgahi H, Grabe HJ, Groenewold NA, Ho TC, Hong SB, Houenou J, Irungu B, Jahanshad N, Khazaie H, Kim H, Koshmanova E, Kocevska D, Kochunov P, Lakbila-Kamal O, Leerssen J, Li M, Luik AI, Muto V, Narbutas J, Nilsonne G, O’Callaghan VS, Olsen A, Osorio RS, Poletti S, Poudel G, Reesen JE, Reneman L, Reyt M, Riemann D, Rosenzweig I, Rostampour M, Saberi A, Schiel J, Schmidt C, Schrantee A, Sciberras E, Silk TJ, Sim K, Smevik H, Soares JC, Spiegelhalder K, Stein DJ, Talwar P, Tamm S, Teresi G l, Valk SL, Someren EV, Vandewalle G, Egroo MV, Völzke H, Walter M, Wassing R, Weber FD, Weihs A, Westlye LT, Wright MJ, Wu M-J, Zak N, Zarei M. 2021. ENIGMA-Sleep: Challenges, opportunities, and the road map. J Sleep Res.

Tahmasian M, Samea F, Khazaie H, Zarei M, Kharabian Masouleh S, Hoffstaedter F, Camilleri J, Kochunov P, Yeo BTT, Eickhoff SB, Valk SL. 2020. The interrelation of sleep and mental and physical health is anchored in grey-matter neuroanatomy and under genetic control. Commun Biol. 3:1–13.

Tai XY, Chen C, Manohar S, Husain M. 2022. Impact of sleep duration on executive function and brain structure. Commun Biol. 5:1–10.

Tashjian SM, Galván A. 2020. Neural recruitment related to threat perception differs as a function of adolescent sleep. Dev Sci. 23:e12933.

Tashjian SM, Goldenberg D, Monti MM, Galván A. 2018. Sleep quality and adolescent default mode network connectivity. Soc Cogn Affect Neurosci. 13:290–299.

Thurman SM, Wasylyshyn N, Roy H, Lieberman G, Garcia JO, Asturias A, Okafor GN, Elliott JC, Giesbrecht B, Grafton ST, Mednick SC, Vettel JM. 2018. Individual differences in compliance and agreement for sleep logs and wrist actigraphy: A longitudinal study of naturalistic sleep in healthy adults. PLOS ONE. 13:e0191883.

Tustison NJ, Avants BB, Cook PA, Zheng Y, Egan A, Yushkevich PA, Gee JC. 2010. N4ITK: Improved N3 Bias Correction. IEEE Trans Med Imaging. 29:1310–1320.

Uy JP, Galván A. 2017. Sleep duration moderates the association between insula activation and risky decisions under stress in adolescents and adults. Neuropsychologia. 95:119–129.

van de Langenberg SCN, Kocevska D, Luik AI. 2022. The multidimensionality of sleep in population-based samples: a narrative review. J Sleep Res. 31.

Vanderhasselt MA, De Raedt R, De Paepe A, Aarts K, Otte G, Van Dorpe J, Pourtois G. 2014. Abnormal proactive and reactive cognitive control during conflict processing in major depression. J Abnorm Psychol. 123:68–80.

Wainberg M, Jones SE, Beaupre LM, Hill SL, Felsky D, Rivas MA, Lim ASP, Ollila HM, Tripathy SJ. 2021. Association of accelerometer-derived sleep measures with lifetime psychiatric diagnoses: A cross-sectional study of 89,205 participants from the UK Biobank. PLOS Med. 18:e1003782.

Walker MP, Stickgold R. 2006. Sleep, Memory, and Plasticity. Annu Rev Psychol. 57:139–166.

Wang Y, Jiang P, Tang S, Lu L, Bu X, Zhang L, Gao Y, Li H, Hu X, Wang S, Jia Z, Roberts N, Huang X, Gong Q. 2021. Left superior temporal sulcus morphometry mediates the impact of anxiety and depressive symptoms on sleep quality in healthy adults. Soc Cogn Affect Neurosci. 16:10.

Watson NF, Badr MS, Belenky G, Bliwise DL, Buxton OM, Buysse D, Dinges DF, Gangwisch J, Grandner MA, Kushida C, Malhotra RK, Martin JL, Patel SR, Quan SF, Tasali E. 2015. Recommended Amount of Sleep for a Healthy Adult: A Joint Consensus Statement of the American Academy of Sleep Medicine and Sleep Research Society. J Clin Sleep Med. 11:591–592.

Wei T, Simko W. 2021. R package “corrplot”: Visualization of a Correlation Matrix.

Wei Y, Van Someren EJ. 2020. Interoception relates to sleep and sleep disorders. Curr Opin Behav Sci, Sleep and cognition. 33:1–7.

Wickham H, Averick M, Bryan J, Chang W, McGowan LD, François R, Grolemund G, Hayes A, Henry L, Hester J, Kuhn M, Pedersen TL, Miller E, Bache SM, Müller K, Ooms J, Robinson D, Seidel DP, Spinu V, Takahashi K, Vaughan D, Wilke C, Woo K, Yutani H. 2019. Welcome to the Tidyverse. J Open Source Softw. 4:1686.

Wild CJ, Nichols ES, Battista ME, Stojanoski B, Owen AM. 2018. Dissociable effects of self-reported daily sleep duration on high-level cognitive abilities. Sleep. 41.

Woo C-W, Krishnan A, Wager TD. 2014. Cluster-extent based thresholding in fMRI analyses: Pitfalls and recommendations. NeuroImage. 91:412–419.

Woolrich MW, Behrens TEJ, Beckmann CF, Jenkinson M, Smith SM. 2004. Multilevel linear modelling for FMRI group analysis using Bayesian inference. NeuroImage. 21:1732–1747.

Woolrich MW, Ripley BD, Brady M, Smith SM. 2001. Temporal autocorrelation in univariate linear modeling of FMRI data. NeuroImage. 14:1370–1386.

Worsley KJ. 2011. Statistical analysis of activation images. In: Jezzard P,, Matthews PM,, Smith SM, editors. Functional Magnetic Resonance Imaging: An Introduction to Methods. Oxford University Press. p. 251–270.

Wylie GR, Pra Sisto AJ, Genova HM, DeLuca J. 2022. Fatigue Across the Lifespan in Men and Women: State vs. Trait. Front Hum Neurosci. 16.

Yaffe K, Nasrallah I, Hoang TD, Lauderdale DS, Knutson KL, Carnethon MR, Launer LJ, Lewis CE, Sidney S. 2016. Sleep Duration and White Matter Quality in Middle-Aged Adults. Sleep. 39:1743–1747.

Yang Y, Miskovich TA, Larson CL. 2018. State Anxiety Impairs Proactive but Enhances Reactive Control. Front Psychol. 9.

Yarkoni T, Barch DM, Gray JR, Conturo TE, Braver TS. 2009. BOLD Correlates of Trial-by-Trial Reaction Time Variability in Gray and White Matter: A Multi-Study fMRI Analysis. PLOS ONE. 4:e4257.

Zhang R, Tomasi D, Shokri-Kojori E, Wiers CE, Wang G-J, Volkow ND. 2020. Sleep inconsistency between weekends and weekdays is associated with changes in brain function during task and rest. Sleep. 43.

Zhang Y, Brady M, Smith S. 2001. Segmentation of brain MR images through a hidden Markov random field model and the expectation-maximization algorithm. IEEE Trans Med Imaging. 20:45–57.

