## Supplementary Material for "Poorer Sleep Health is Associated With Altered Brain Activation During Cognitive Control Processing in Healthy Adults"

**Supplementary Table S1** *fMRI Task Performance and Self-Report Measures (N = 81)*

| Variable | Mean / SD | Min | Max |
|---|---|---|---|
| Hit RT, mean (ms) | 396.48 (43.36) | 312.40 | 573.43 |
| Δ Hit RT, mean (ms) | 1.66 (17.95) | -39.98 | 74.15 |
| Hit RT SD, mean (ms) | 64.08 (17.36) | 39.76 | 129.94 |
| Δ Hit RT SD, mean (ms) | 0.54 (17.86) | -30.18 | 89.30 |
| Omission errors, sum (n) | 0.79 (1.41) | 0 | 6 |
| Δ Omission errors, sum (n) | 0.25 (1.08) | -2 | 6 |
| Commission errors, sum (n) | 17.62 (10.14) | 1 | 41 |
| Δ Commission errors, sum (n) | 0.69 (2.00) | -6 | 5 |
| Detectability (d') | 3.27 (0.71) | 1.68 | 4.92 |
| Δ Detectability (d') | -0.22 (0.60) | -1.56 | 1.27 |
| Fatigue during task (1-10)[a] | 4.63 (1.65) | 1 | 9 |
| Sleepiness during task (1-10)[a] | 4.81 (1.55) | 1 | 8 |

Δ indicates time-on-task changes (time epoch 4 - time epoch 1). [a]Self-report measures during task performance were obtained by asking the participants to rate their level of mental fatigue and sleepiness via the MRI speaker system between task runs. Levels of sleepiness were rated according to the Karolinska Sleepiness Scale (Akerstedt and Gillberg 1990). RT = reaction time, ms = milliseconds, SD = standard deviation.

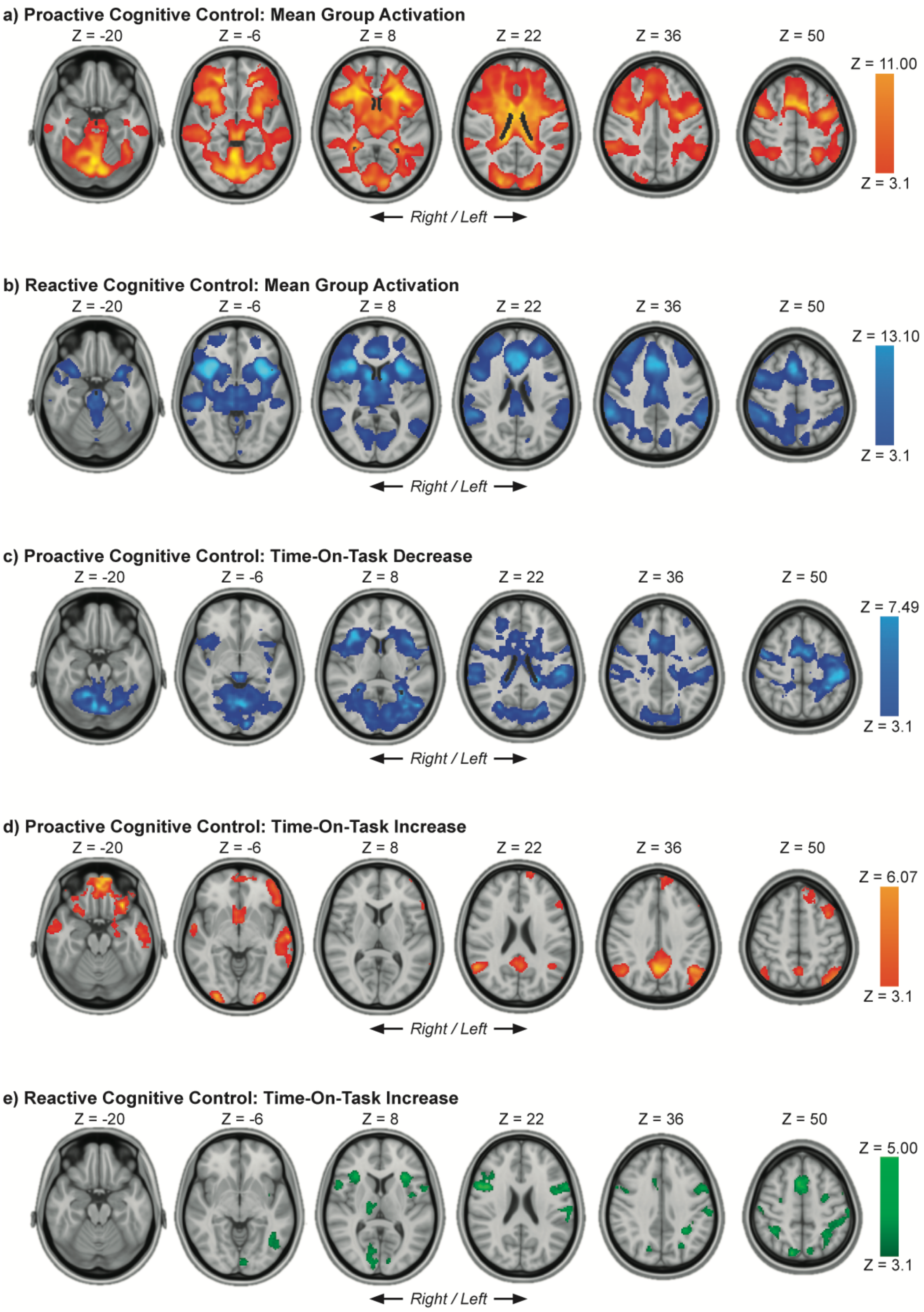


**Supplementary Figure S1. Group Average Cognitive Control Activations.** a) Proactive Cognitive Control, b) Reactive Cognitive Control, c) Proactive Cognitive Control TOT Decrease, d) Proactive Cognitive Control TOT Increase, e) Reactive Cognitive Control TOT Increase. Results were obtained using mixed-effects models and are presented on a 1-mm MNI standard space template. Cluster-based inference was used to control the family-wise error rate in each model (cluster-defining threshold = $Z > 3.1$, cluster probability threshold = $p < .05$). Due to strong activation in the main contrasts (figure a, b), resulting clusters are very large and span over several brain regions. Slices that are most representative for the overall findings (anatomical, and across different clusters) have been selected. As these are 2D representations of 3D volumes, some of the clusters may only be partly visible (see Supplementary Tables S2, S3 for details on cluster size/coordinates). TOT = time on task, MNI = Montreal Neurological Institute.

**Supplementary Table S2** *Group Average Activations for Proactive and Reactive Cognitive Control Processing*

| **Contrast / Region** | **Right / Left** | **Size (#voxels)** | **P Value** | **Peak Z Value** | **Peak Coordinates (MNI)** | | |
|---|---|---|---|---|---|---|---|
| | | | | | **X** | **Y** | **Z** |
| **Proactive Cognitive Control** | *Minimum significant cluster size = 183 voxels* | | | | | | |
| Insular Cortex / Frontal Operculum / Frontal Orbital Cortex | R / L | 93854 | < .0001 | 11.00 | -24 | 20 | 12 |
| Frontal Operculum | L | lm | - | 11.00 | -38 | 16 | 14 |
| Insular Cortex / Frontal Operculum | R | lm | - | 10.90 | 26 | 22 | 12 |
| Lingual Gyrus | R | lm | - | 10.80 | 8 | -76 | -16 |
| Cerebellum – Right V | R | lm | - | 10.80 | 4 | -58 | -8 |
| Lingual Gyrus / Cerebellum – Left VI | L | lm | - | 10.70 | -6 | -76 | -16 |
| **Reactive Cognitive Control** | *Minimum significant cluster size = 170 voxels* | | | | | | |
| Insular Cortex | R / L | 77594 | < .0001 | 13.10 | -36 | 16 | -6 |
| Insular Cortex | R | lm | - | 13.10 | 38 | 16 | -4 |
| Insular Cortex / Frontal Operculum | R | lm | - | 12.90 | 34 | 22 | 2 |
| Insular Cortex | L | lm | - | 12.80 | -30 | 22 | -2 |
| Insular Cortex / Frontal Orbital Cortex | R | lm | - | 12.60 | 32 | 20 | -8 |
| Paracingulate Gyrus | R | lm | - | 11.50 | 4 | 22 | 42 |
| Cerebellum – Right Crus I | R | 367 | .0012 | 5.73 | 38 | -52 | -32 |

Main peaks and local maxima (lm) within each cluster are reported. Results were obtained using mixed-effects models and cluster-based inference was used to control the family-wise error rate in each model (cluster-defining threshold = Z > 3.1, cluster probability threshold = p < .05). *Minimum significant cluster size* indicates the minimum number of contiguous voxels (Z > 3.1) required for a cluster to be considered significant (p <. 05), determined using Gaussian RFT. Anatomical regions for each peak Z value were labeled using visual inspection and the Harvard Oxford cortical / subcortical structural atlases and the Cerebellar atlas (normalized with FLIRT) within the FSL software. Due to very strong activations in these main contrasts, the clusters are large and span over several brain regions (see Supplementary Figure S1 for visualization). RFT = Random Field Theory, MNI = Montreal Neurological Institute, R = Right, L = Left.

**Supplementary Table S3** *Group Average Time-On-Task Changes for Proactive and Reactive Cognitive Control Processing*

| Contrast / Region | Right / Left | Size (#voxels) | P Value | Peak Z Value | Peak Coordinates (MNI) | | |
|---|---|---|---|---|---|---|---|
| | | | | | X | Y | Z |
| **Proactive Cognitive Control:** | *Minimum significant cluster size = 187 voxels* | | | | | | |
| **Time-On-Task Decrease** | | | | | | | |
| Superior Parietal Lobule / Postcentral Gyrus | R / L | 61822 | < .0001 | -7.49 | -32 | -46 | 58 |
| Juxtapositional Lobule | R | lm | - | -7.33 | 12 | 2 | 56 |
| Cerebellum – Right V / VI | R | lm | - | -7.24 | 6 | -64 | -14 |
| Left Lateral Ventricle | L | lm | - | -7.20 | -16 | -30 | 18 |
| Lingual Gyrus / Cerebellum | R | lm | - | -7.17 | 8 | -72 | -14 |
| Superior Parietal Lobule / Postcentral Gyrus | L | lm | - | -7.09 | -42 | -40 | 60 |
| **Proactive Cognitive Control:** | *Minimum significant cluster size = 187 voxels* | | | | | | |
| **Time-On-Task Increase** | | | | | | | |
| Frontal Orbital Cortex | L | 4938 | < .0001 | 6.07 | -30 | 26 | -18 |
| Middle Temporal Gyrus | L | 2654 | < .0001 | 5.52 | -58 | -14 | -14 |
| Lateral Occipital Cortex | L | 2166 | < .0001 | 5.63 | -44 | -66 | 56 |
| Precuneus Cortex | R | 2138 | < .0001 | 5.51 | 4 | -58 | 34 |
| Suprior Temporal Gyrus | R | 1516 | < .0001 | 4.71 | 60 | -4 | -8 |
| Angular Gyrus | R | 1336 | < .0001 | 5.59 | 52 | -56 | 26 |
| Cerebellum: Right Crus I | R | 737 | < .0001 | 4.86 | 22 | -82 | -28 |
| Cerebellum – Left Crus II | L | 634 | < .0001 | 5.09 | -18 | -84 | -38 |
| Occipital Pole | R | 338 | .0036 | 5.27 | 28 | -96 | -4 |
| Occipital Pole | L | 300 | .0066 | 5.80 | -26 | -96 | -10 |
| **Reactive Cognitive Control:** | *Minimum significant cluster size = 168 voxels* | | | | | | |
| **Time-On-Task Increase** | | | | | | | |
| Superior Parietal Lobule | L | 1791 | < .0001 | 4.23 | -26 | -48 | 40 |
| Inferior Frontal Gyrus, pars opercularis | L | 1658 | < .0001 | 4.55 | -58 | 12 | 26 |
| Inferior Frontal Gyrus, pars opercularis | R | 1271 | < .0001 | 5.00 | 54 | 12 | 18 |
| Paracingulate Gyrus | R | 984 | < .0001 | 4.59 | 6 | 10 | 52 |
| Occipital Pole | R | 434 | .0003 | 4.05 | 8 | -90 | 4 |
| Inferior Temporal Gyrus | L | 433 | .0003 | 4.62 | -42 | -50 | -10 |
| Precentral Gyrus | R | 407 | .0005 | 4.00 | 46 | -8 | 60 |

**Supplementary Table S3 Continued**

| Contrast / Region | Right / Left | Size (#voxels) | P Value | Peak Z Value | Peak Coordinates (MNI) | | |
|---|---|---|---|---|---|---|---|
| | | | | | X | Y | Z |
| Superior Parietal Lobule | R | 404 | .0006 | 4.03 | 38 | -48 | 50 |
| Lateral Occipital Cortex, superior division | R | 282 | .0049 | 4.56 | 14 | -78 | 48 |
| Postcentral Gyrus | L | 205 | .0223 | 3.88 | -46 | -16 | 58 |
| Precuneus Cortex | R | 197 | .0263 | 4.18 | -8 | -72 | 46 |
| Precentral Gyrus | L | 186 | .0332 | 3.82 | -26 | -8 | 54 |
| Thalamus | R | 186 | .0332 | 4.40 | 12 | -16 | 6 |
| Postcentral Gyrus | R | 185 | .0339 | 3.73 | 54 | -20 | 42 |

Main peaks within each cluster are reported, and for Proactive Cognitive Control Decrease also local maxima (lm) are included due to the cluster's size and span. Results were obtained using mixed-effects models and cluster-based inference was used to control the family-wise error rate in each model (cluster-defining threshold = $Z > 3.1$, cluster probability threshold = $p < .05$). *Minimum significant cluster size* indicates the minimum number of contiguous voxels ($Z > 3.1$) required for a cluster to be considered significant ($p <. 05$), determined using Gaussian RFT. Anatomical regions for each peak Z value were labeled using visual inspection and the Harvard Oxford cortical / subcortical structural atlases and the Cerebellar atlas (normalized with FLIRT) within the FSL software. Note that some clusters are relatively large and therefore span over several brain regions (see Supplementary Figure S1 for visualization). RFT = Random Field Theory, MNI = Montreal Neurological Institute, R = Right, L = Left.
